# Succinate and its carrier Sfc1 mediate metabolic control of mitochondrial protein import by the TIM23 translocase

**DOI:** 10.64898/2025.12.08.692925

**Authors:** Koyeli Das, Rituparna Samanta, Tamar Ziv, Johannes M. Herrmann, Bella Kalderon, Ophry Pines

**Affiliations:** Department of Microbiology and Molecular Genetics, The Institute for Medical Research Israel Canada (IMRIC), Faculty of Medicine, The Hebrew University of Jerusalem, P.O.B 12272, Jerusalem 91120, Israel; Department of Chemical, Biological and Materials Engineering, University of South Florida, Tampa, Florida 33620; The Smoler Proteomics Center, Technion Israel Institute of Technology, Haifa, Israel; Cell Biology, University of Kaiserslautern, RPTU, Erwin-Schrödinger-Strasse 13, 67663 Kaiserslautern, Germany

**Keywords:** mitochondrial protein import, metabolic signaling, tricarboxylic acid cycle, glyoxylate shunt, TIM23 complex, succinate-fumarate carrier Sfc1, dual targeting, aconitase, fumarase, metabolites

## Abstract

Tim23 is an essential component of the mitochondrial inner membrane translocase and Sfc1 is a carrier that exchanges succinate for fumarate across that membrane. Sfc1 and succinic acid availability regulate dual targeting of fumarase and aconitase by facilitating mitochondrial import of their newly synthesized precursors, as shown by pulse-chase experiments. Here we show that Sfc1 directly interacts with the Tim23 and succinate affects this interaction, which in turn affects mitochondrial protein import. Physical interaction between Tim23 and Sfc1 was proven by co immunoprecipitation, Bimolecular Fluorescence Complementation (BiFC) and Biotin-based proximity labeling (TurboID). Proximity labeling-informed mutagenesis allowed us to dissect the carrier activity of Sfc1 from its function as a TIM23 regulator. We performed Rosetta-MP docking of Sfc1 and Tim23 to envisage the interface. Thus, our findings show that metabolites can regulate mitochondrial import and adjust the segregation of key metabolic enzymes between the cytosol and mitochondria.

## Introduction

Fumarase and aconitase in yeast are dual localized to the cytosol and mitochondria by a similar mechanism. These two tricarboxylic acid cycle enzymes are single translation products that are targeted to and processed by mitochondrial processing peptidase (MPP) in mitochondria, prior to distribution. The mechanism includes reverse translocation of a subset of processed molecules back into the cytosol ^1–8^. We have previously shown that depletion of enzymes of the glyoxylate shunt (deletion mutations Δ*cit2*, Δ*mls1*, Δ*aco1* and Δ*icl1*) alters fumarase dual distribution, causing the vast majority of fumarase to be fully imported into mitochondria with a small amount or no fumarase in the cytosol ^9^. Thus, dual distribution of fumarase (similar amounts in the cytosol and mitochondria) depends on an active glyoxylate shunt. Intriguingly, when succinic acid, a product of the glyoxylate shunt, is added to the growth medium, fumarase dual distribution is altered such that there are lower levels of fumarase in the cytosol ^9^.

Aconitase, like fumarase is a TCA cycle enzyme, yet it is also a component of the glyoxylate shunt in the yeast cytosol, a pathway which is required for growth on non-fermentable carbon sources ^10^. Succinate is a product of the glyoxylate shunt and aconitase is a dual targeted protein with a very small fraction (<5%) in the cytosol, a situation termed “eclipsed distribution” ^10,11^. The C-terminal domain of aconitase drives dual localization by its interaction with the N-terminal and thereby affects mitochondrial import ^4,12^.

Mitochondrial carriers facilitate the transport of various molecules across the inner membrane, connecting cytoplasmic and matrix pathways to support cellular function ^13–15^. Sfc1 (Succinate Fumarate Carrier 1) ^16–18^ facilitates exchange of succinate for fumarate by a strict counter-exchange mechanism (Figure 1A, schematic illustration). The exported fumarate via the TCA cycle can participate, for example, in the gluconeogenic pathway in the cytosol ^19,20^. This allows *S. cerevisiae* growth on ethanol or acetate as a single carbon source ^16,21–23^. Sfc1 is, in fact, a component of the glyoxylate shunt and thus a Sfc1 knockout strain (Δ*sfc1*) cannot grow on ethanol and/or acetate as a sole carbon source (Figure 1C).

**Figure 1.**
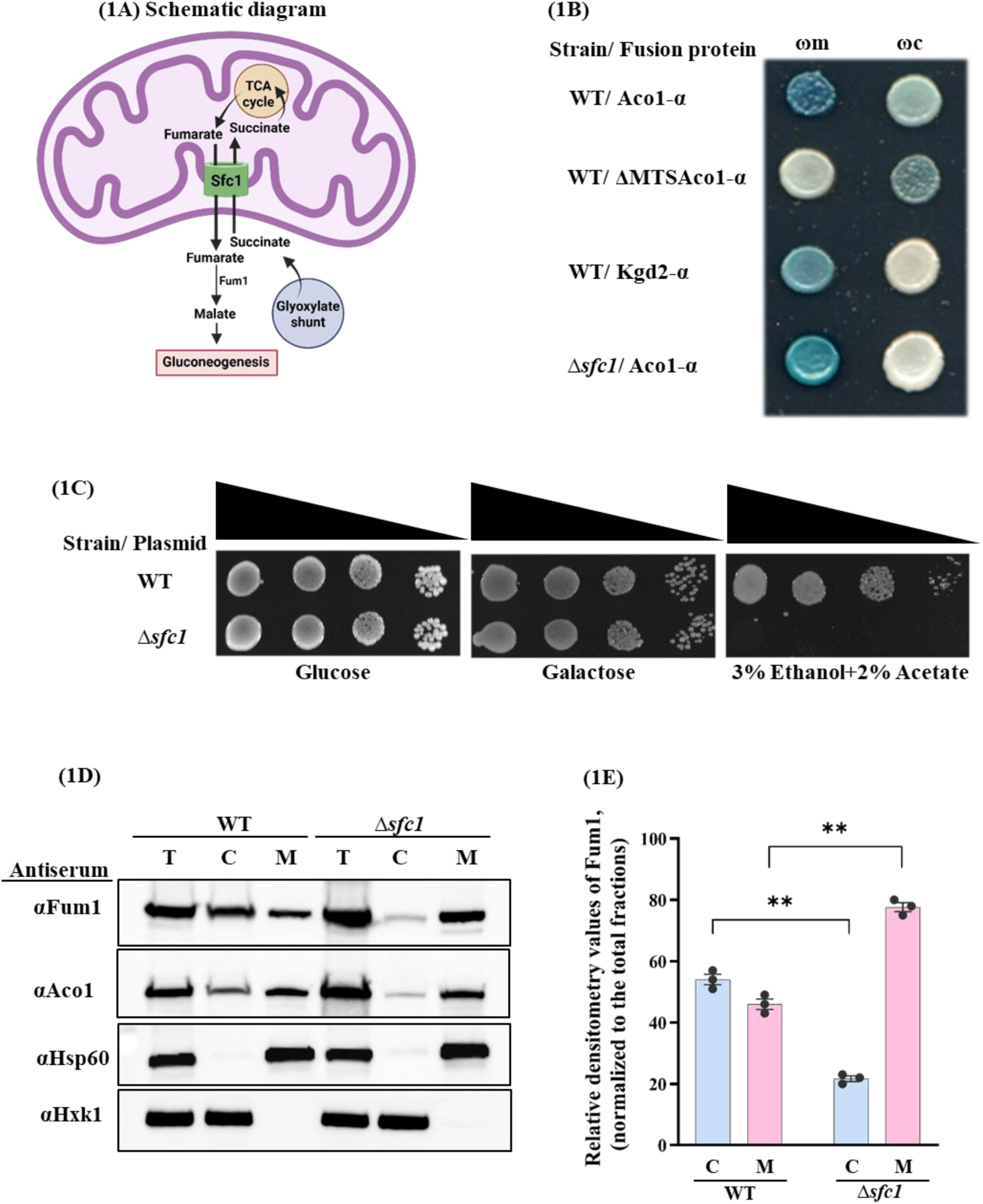
Sfc1 determines the distribution of aconitase and fumarase. **(1A)** Schematic diagram of Succinate-fumarate carrier (Sfc1). Sfc1 transports succinate, produced in the cytosol by the glyoxylate shunt, into the mitochondrial matrix in exchange for the export of fumarate from mitochondria to the cytosol. Succinate in mitochondria can enter the TCA cycle while fumarate exported by Sfc1 can be utilized by the gluconeogenic pathway. **(1B)** Yeast cultures expressing either cytosolic ω (ωc) or mitochondrial ω (ωm) together with Aco1-α fused proteins were grown on galactose media supplemented with X-gal. Aco1-α, which is eclipsed dual distributed between mitochondria and the cytosol, exhibits blue colonies with both ωc and ωm in wild-type cells (WT), top row. The same Aco1-α is exclusively localized to mitochondria in Δ*sfc1* cells (lacking a functional Sfc1 protein), bottom row. Controls include ΔMTSAco1-α (Aconitase lacking the mitochondrial targeting signal) and Kgd2-α (Dihydrolipoyl transsuccinylase 2), acting as cytosolic and mitochondrial markers, second and third rows respectively. **(1C)** Wild-type and Sfc1-KO strains were diluted and grown on glucose, galactose and ethanolacetate plates. Δ*sfc1* cannot grow on ethanol-acetate as a sole carbon source since Sfc1 is required for the glyoxylate shunt. **(1D)** Wild-type (WT) and Δ*sfc1* yeast strains were grown in galactose medium and subjected to subcellular fractionation. Equivalent aliquots of the total (T) cytosolic (C) and mitochondrial (M) fractions were analyzed by western blotting using antibodies against the indicated proteins. **(1E)** The percentages of fumarase in the cytosolic (C) and mitochondrial (M) fractions was calculated based on densitometry analysis of band intensities and normalized to the total (T) fractions. The graph represents the quantification from three independent experiments (mean ± SEM [n=3], two-tailed student’s t-test cytosolic **p=0.004 and mitochondrial **p=0.007).

Here we show that the cytosolic versus mitochondrial distribution of a protein is governed by intracellular metabolite cues, in this case, the direct interaction of Sfc1 (succinate-fumarate exchange carrier) with Tim23 (a component of the inner membrane translocase). The intriguing question is how metabolites of the glyoxylate shunt can control the distribution of proteins in different compartments of eukaryotic cells. In fact, the results shown, for first time, in this study, identify metabolites, components and mechanisms of an important regulatory circuit that can control the distribution of proteins in the different compartments of eukaryotic cells.

## Results

### Sfc1 affects the dual targeting of certain mitochondrial proteins

Sfc1 is a mitochondrial inner membrane succinate-fumarate exchange carrier which imports succinate into mitochondria in exchange for fumarate (Figure1A). Depletion of Sfc1 (Δ*sfc1*) similarly to the depletion of enzymes of the glyoxylate shunt (e.g. Δ*cit2, Δmls1 and Δicl1*) leads to inability to grow on ethanol and/or acetate as a sole carbon source ^9^ (Figure 1C). We employed the split gene α-complementation assay ^24–28^ to identify the subcellular localization of aconitase in a Sfc1 deletion strain (termed Δ*sfc1,* Aco1-α, Figure 1B). The α-complementation assay utilizes α (77 amino acids) and ω (993 amino acids) fragments of β-galactosidase, which assemble *in vivo* to form an active enzyme complex. This method, prevents a predominant protein (in a different compartment) from generating a signal on its own, and overshadowing an eclipsed protein’s signal. A schematic representation of this approach is shown in Figure S1. To investigate the potential role of Sfc1 in the subcellular distribution of aconitase, we constructed a Δ*sfc1* strain lacking a functional

*SFC1* gene and expressing the α-fragment of β-galactosidase fused to the C-terminus of aconitase (Δ*sfc1,* Aco1-α). Aco1-α was co-expressed in the Δ*sfc1* strain with either cytosolic ωc or mitochondrial ωm fragments of β-galactosidase and grown on galactose plates containing X-gal (Figure 1B). Blue colonies formed by Δ*sfc1* cells co-expressing Aco1-α with ωm and not with ωc, indicate that it is exclusively mitochondrial (Figure 1B, bottom row). This is in contrast to WT expressing the Aco1-α which produces blue colonies with both ωm and ωc, thus, demonstrating dual localization (Figure 1B, top row). Accordingly, ΔMTS-Aco1-α (Aconitase lacking the mitochondrial targeting signal) is a cytosolic marker and produced blue colonies only when co expressed with ωc (Figure 1B, second row). As an additional control, Kgd2 (Dihydrolipoyl transsuccinylase 2), a specific mitochondrial marker fused to the α-fragment, produces blue colonies only when co-expressed with ωm (Figure 1B, third row).

Regarding fumarase, yeast wild-type (WT) and Sfc1 knock out (KO) strains were grown in galactose medium and subjected to subcellular fractionation (Figure 1D). Equal aliquots of total (T), cytosolic (C), and mitochondrial (M) fractions of fumarase in both strains were analyzed by western blot analysis using an anti-fumarase antibody (Top panel). The distribution of fumarase is altered in the Δ*sfc1* strain, in such a way that there are lower levels of fumarase in the cytosol and higher levels of fumarase in mitochondria and this was quantified using densitometry analysis of band intensities. We observed that in Δ*sfc1*, the cytosolic amount of fumarase decreased from 54% to 22%, while the mitochondrial fraction increased from 46% to 78% compared to WT (Figure 1E). This indicates that deletion of Sfc1 promotes the import of these proteins into mitochondria and suggests that Sfc1, modulates the fumarase and aconitase distribution between mitochondria and the cytosol (Figure 1B).

### Sfc1 and its substrate, succinate, affect the rate of protein import into mitochondria

Pulse-chase labeling experiments of yeast aconitase (Aco1) were undertaken. Our experimental approach involves inhibiting mitochondrial protein import *in vivo* using CCCP, while simultaneously labeling yeast cells with ^35^S-methionine to track Aco1. Subsequently, we reverse the import block using DTT. Samples are then collected at specified intervals, and labeled proteins are immunoprecipitated and subjected to SDS-PAGE followed by autoradiography (Figure S2, schematic diagram). Using this approach, the fully translated precursor is accumulated in the cytosol and only then (when we remove the translocation block with DTT) is it translocated post translationally. This allowed us to examine the rate of mature aconitase appearance (processed) as an indication of translocation. Importantly, as shown in Figure 2A, addition of 10mM mono-ethyl succinate induced a significant increase in the Aco1 import rate into mitochondria, as compared to control cells, or cells incubated in the presence of 10mM di-ethyl malate (Figures 2A and 2B). To ask whether the intracellular levels of Sfc1 can affect Aco1 mitochondrial import, WT cells, Sfc1 deleted (Δ*sfc1*) and cells transformed with a plasmid over-expressing Sfc1 (pSfc1), were examined. It turns out that knockout of Sfc1 (Δ*sfc1*), facilitates aconitase import into mitochondria as compared to the WT (Figures 2C and 2D). Δ*sfc1* cells incubated with succinic acid, exhibited no additional effect on aconitase import (Figures 2C and 2D). Accordingly, when Sfc1 is overexpressed (pSfc1), aconitase import into mitochondria is reduced (Figures 2E and 2F) again suggesting that Sfc1 can inhibit the efficient protein translocation into mitochondria. Therefore, we conclude that Sfc1 controls fumarase and aconitase import by reducing translocation rates; the presence of the Sfc1 substrate succinate, apparently alleviates the Sfc1-mediated inhibition.

**Figure 2.**
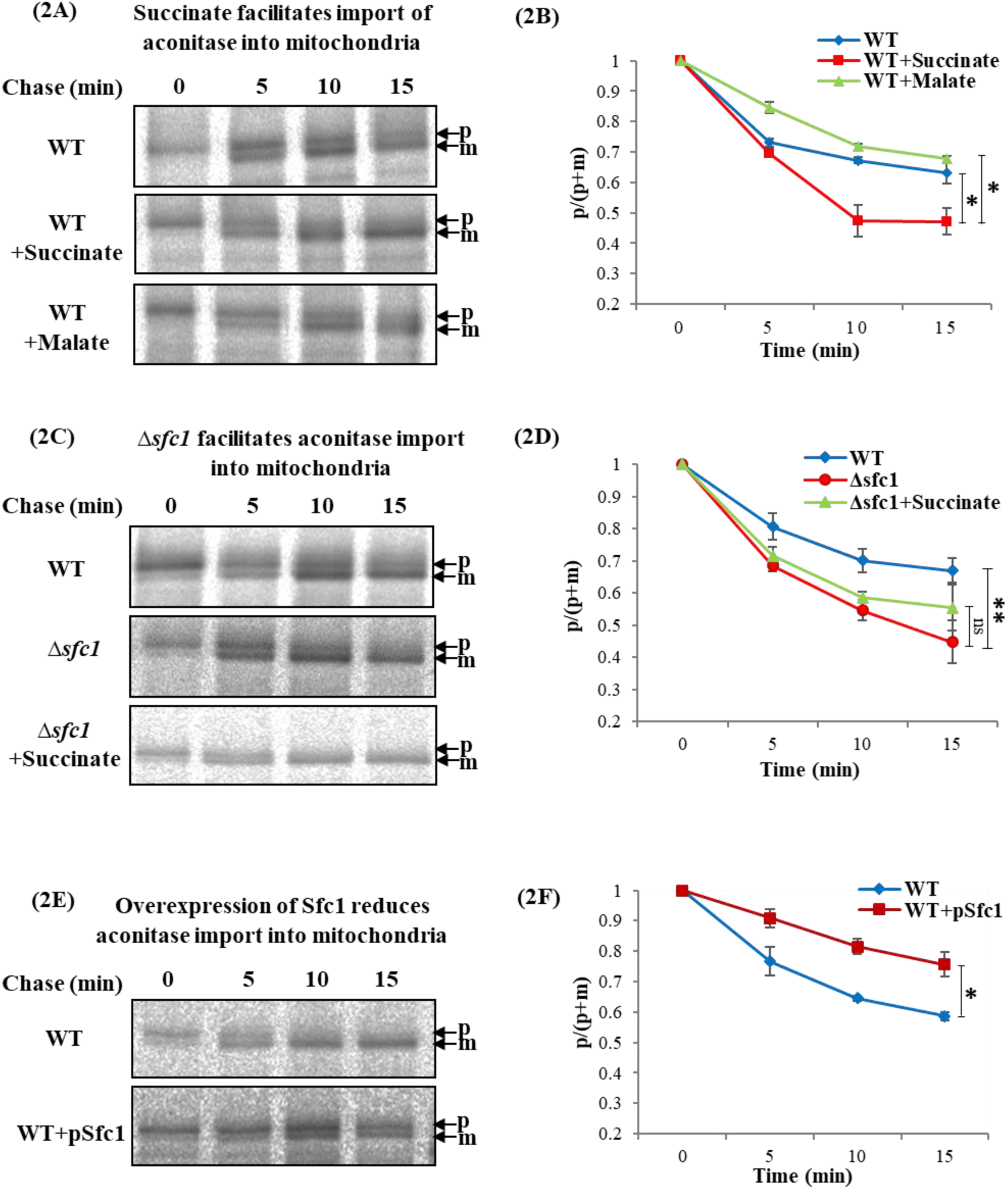
Sfc1 influences the efficiency of protein import into mitochondria. The effect of succinate **(2A)**, Δ*sfc1* **(2C)** and Sfc1 overexpression **(2E)** on aconitase (Aco1) import by pulse-chase experiments *in vivo*. The left panels (2A, 2C, and 2E) show [^35^S] autoradiography while the right panels (2B, 2D, and 2F) present quantification. p−precursor (with the mitochondrial targeting signal, MTS), m−mature (without the MTS) and p/(p+m) is a calculated parameter indicating the efficiency of mitochondrial import. The arrows indicate the positions of precursor (p) and mature (m) proteins. With time precursor (p) is translocated into mitochondria and the mature fraction (m) increases (at the expense of the precursor). Precursor (p) and mature (m) band intensities were quantified by densitometry and normalized to the precursor (p) intensity at 0min (set to 1) for each treatment. Each graph represents three independent experiments (mean ± SEM [n=3]. Statistical significance was assessed using student’s t-test: **(2B)** WT vs. WT+Succinate (*p< 0.05) and WT+Succinate vs. WT+Malate (*p< 0.05) **(2D)** WT vs. Δ*sfc1* (\**\**p<0.01) and **(2F)** WT vs. WT+pSfc1 (*\**p< 0.05).

To further support the results obtained in intact cells, experiments were undertaken with isolated mitochondria, *in vitro*. Isolated mitochondria were incubated with ^35^S-methionine radiolabeled Aco1 precursors (Figure S3A, schematic illustration). Samples were withdrawn at different time points and further incubated in the presence of proteinase K (PK) to digest the proteins on the mitochondrial surface, so that only completely imported ^35^S-labeled Aco1 molecules (mature proteins) are detected. We observe that the rate of Aco1 uptake into isolated mitochondria (Figure S3B, left panel gel and right panel graph) is facilitated by succinate when compared to its import in the absence of acid or in the presence of malate. The overall effect of succinate treatment was smaller than that observed *in vivo* (Figure 2), yet was also statistically significant. Previous studies in our lab demonstrated that succinate does not influence the *in vitro* import of other mitochobdrial proteins including Su9-DHFR and F1β into isolated mitochondria ^9^.

The expression of Sfc1 across different genetic backgrounds was also considered and as expected yeast extracts analyzed by western blot revealed that overexpression of Sfc1 (by pSfc1) leads to approximately 3.4-fold higher expression than in the WT (Figure S4, compare lanes 7-9 to lanes 1-3), while Sfc1 in a Δ*sfc1* knockout strain is undetectable (Figure S4, lanes 4-6). Worth pointing out is that succinate levels do not affect Aco1 or Sfc1 expression (compare each triplet of lanes to itself).

### Do Tim23 and Sfc1 physically interact?

Based on the observations above, we hypothesized the existence of a physical interaction between Tim23 and Sfc1. Accordingly, we have addressed the question above by multiple approaches as specified below. This was initially examined by co-immunoprecipitation (Co-IP) of Sfc1 from cell extracts, which was subsequently analysed by western blotting with anti-Sfc1 antibody (Figure 3A, top panel) and anti-Tim23 antibody (Figure 3A, bottom panel). The results revealed that Tim23 was co-immunoprecipitated with Sfc1 (Figure 3A, lane 1, indicated by a star), suggesting a physical interaction between these two proteins, which is obviously absent in the Δ*sfc1* strain (Figure 3A lane 2).

**Figure 3.**
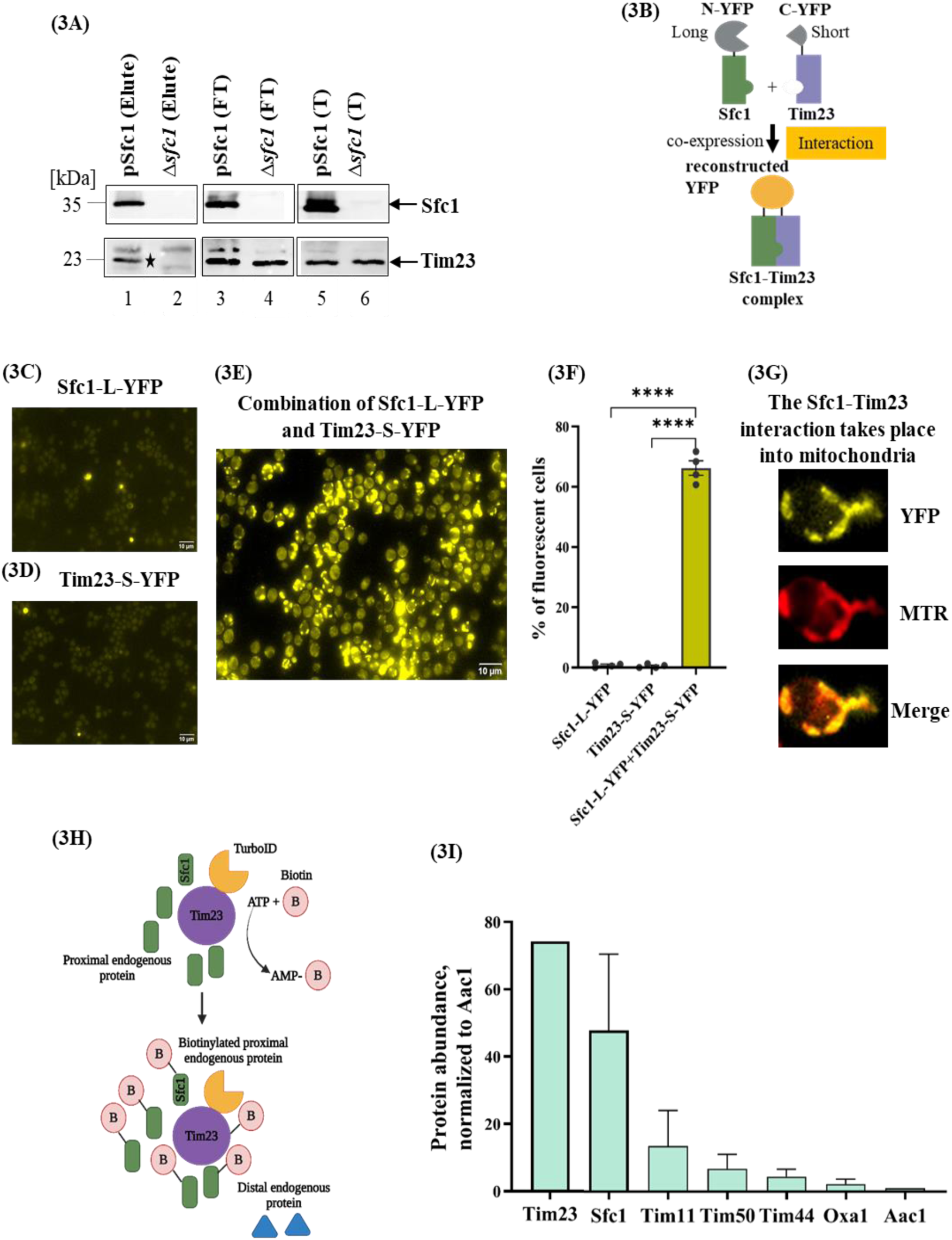
Co-immunoprecipitation (Co-IP) of Tim23 with Sfc1. **(3A)** Cell lysates were immunoprecipitated with anti-Sfc1 antibody and subsequently analyzed by SDS-PAGE and western blot with anti-Sfc1 (top panels) or anti-Tim23 (bottom panels) antibodies. Lanes 1-2, 34, 5-6 represent elute (E), flow-through (FT), and total (T) fractions of pSfc1 and Δ*sfc1* strains respectively. In lane 1, the band indicated by a star represents the co-immunoprecipitated Tim23. **Interactions of Sfc1 and Tim23 analyzed by Bimolecular Fluorescence Complementation (BiFC) *in vivo*.** Schematic illustration of BiFC technology: Sfc1 and Tim23 were tagged with YFP fragments and interaction between Sfc1 and Tim23 was followed by YFP fluorescence **(3B).** Yeast cells transformed either with a plasmid encoding Sfc1 fused to long YFP **(3C)** or a plasmid encoding Tim23 fused to short YFP **(3D),** show insignificant fluorescence (the bright fluorescent cells are dead cells). Cells transformed with both Sfc1 fused to long YFP and Tim23 fused to short YFP **(3E)**, display highly significant fluorescence. **(3F)** The quantification in the graph represents the percentage of fluorescent cells out of the total cell count obtained from four independent experiments (mean ± SEM [n=4], two-tailed student’s t-test ****p<0.0001). **(3G)** Cells expressing Sfc1-long-YFP together with Tim23-short-YFP were incubated with mitotracker red (MTR). The yellow fluorescence merged with red fluorescence indicates direct Sfc1-Tim23 interaction which takes place in the mitochondria. **TurboID identifies Tim23 and Sfc1 as proximate proteins in yeast cells. (3H)** Diagrammatic representation of TurboID-based proximity labeling to identify Tim23 interactors: To identify proteins proximal to Tim23 in yeast, TurboID, a promiscuous biotin ligase was fused to the Nterminus of Tim23. In the presence of ATP and exogenous biotin, TurboID generates biotin-5′AMP, which covalently tags nearby lysine residues within ∼10 nm. **(3I)** Tim23 was fused to the N-terminus of the TurboID biotin ligase, and biotinylated proteins identified by LC-MS/MS are plotted as protein abundance (peptide intensity), normalized to the control protein Aac1. The graph is based on three independent LC-MS/MS experiments, in which Tim23 undergoes selfbiotinylation due to its fusion with TurboID.

To further support the concept that Sfc1 and Tim23 physically interact with each other, we employed an additional approach termed Bimolecular Fluorescence Complementation (BiFC) or split YFP technology ^29,30^. BiFC employs fragments of yellow fluorescent protein (YFP) as molecular tags attached to proteins (Sfc1 and Tim23 in this study). Fluorescence is observed only when these non-fluorescent fragments of YFP are brought into proximity and this can occur due to interactions between the molecules to which they are attached (Figure 3B, illustration). *SFC1* and *TIM23* genes were cloned into yeast expression plasmids under an ADH1 promoter, and were tagged by the appropriate YFP fragments. The expression of Sfc1 and Tim23 tagged with YFP fragments is shown in Figures S5A and S5B respectively.

Yeast cells (*S. cerevisiae*) transformed with a plasmid encoding Sfc1 fused to the long YFP (Figure 3C) or a plasmid encoding Tim23 fused to the short YFP (Figure 3D) do not display fluorescence. In contrast, yeast cells transformed with both plasmids, encoding Sfc1 fused to long YFP and Tim23 fused to the short YFP (Figure 3E) display significant fluorescence, indicating that an interaction between Sfc1 and Tim23, takes place *in vivo*. We quantified the percentage of fluorescent cells (Figure 3F) and found that co-expression of Sfc1 fused to long YFP and Tim23 fused to short YFP displayed ∼66% of fluorescent cells. In contrast, control strains expressing either Sfc1 fused to long YFP or Tim23 fused to short YFP exhibited only 0.8% and 0.5% fluorescent cells, respectively. The punctate appearance of the fluorescence in cells is characteristic of mitochondria. Mitochondria can be labeled *in vivo* with MitoTracker Red CMXRos (MTR) and we found an overlap between the “Sfc1-Tim23” YFP fluorescence and the MTR labeled mitochondrial fluorescence (yellow-orange fluorescence, Figure 3G). This result indicates that a physical interaction between Sfc1 and Tim23 proteins occurs in mitochondria.

A series of control experiments were undertaken. Yeast cells co-transformed with plasmids encoding short YFP and long YFP under a ADH1 promoter, do not exhibit any fluorescence (Figure S5D). This indicates that the reconstitution of the YFP fragments occurs when in close proximity, via the Sfc1 and Tim23 interaction (Figure S5C). To test whether the interaction between Sfc1 and Tim23 is specific, we examined, as controls, three different inner mitochondrial membrane carriers with molecular weights and membrane orientations similar to that of Sfc1; Mme1 (mitochondrial magnesium exporter), Crc1 (carnitine carrier) and Oac1 (oxaloacetate carrier) which were tagged with a split YFP sequence. The expression of short YFP fusions of these three proteins was validated by western blot using an anti-YFP antibody (Figure S6A). Cells expressing Mme1, Crc1 or Oac1 fused to short YFP and Sfc1 fused to long YFP displayed insignificant fluorescence (Figures S6C-S6E) compared to cells expressing Sfc1 fused to the long YFP and Tim23 fused to the short YFP (Figure S6B) which displayed significant fluorescence.

In a parallel set of experiments, yeast cells were transformed with plasmids encoding Tim23 fused to long YFP, along with either Mme1 (Figure S6G), Crc1 (Figure S6H), or Oac1 (Figure S6I) fused to short YFP. The resulting cellular fluorescence of these was insignificant when compared to Tim23 fused to the long YFP and Sfc1 to short YFP (Figure S6F), which showed a significant level of fluorescence. These results indicate that these membrane proteins do not undergo nonspecific interactions within the membrane, and provides support for a specific interaction between Sfc1 and Tim23 in the mitochondrial membrane.

### Proximal biotinylation of Sfc1 by Tim23 fused to TurboID

To further validate the interactions of Sfc1-Tim23, we employed enzyme-catalyzed proximity labeling (PL) mediated by TurboID. TurboID is a promiscuous mutant of the *Escherichia coli* biotin ligase (BirA), which in this study is fused to our protein of interest, Tim23. When biotin is added, it uses ATP to convert biotin into biotin-5’-AMP anhydride, a reactive intermediate that covalently labels proteins within a few nanometers of the enzyme. The biotinylated proteins are then collected using streptavidin-coated beads and identified via mass spectrometry (MS) (Figure 3H, schematic diagram).

To employ TurboID in yeast, we cloned the TIM23 ORF tagged with a sequence encoding TurboID and a V5 tag at the C-terminus (from Addgene). Western blot analysis confirmed the successful cloning and expression of the Tim23 fusion. The tagged Tim23 fusion protein exhibited a molecular mass of approximately 59 kDa, which is about 35 kDa higher than the molecular mass of the untagged Tim23 (23 kDa) (Figure S7A). Worth mentioning is that tagging a bait protein with a biotin ligase derivative, is known to result in significant self-biotinylation of the bait protein ^31,32^. For the sake of brevity, the TurboID-based biotinylation approach in yeast and the description of Tim23 TurboID-V5 expression under different conditions is provided in Figures S7B and S7C, respectively.

To identify proximal proteins and potential physical interaction partners of Tim23, enrichment using streptavidin beads was performed. The biotinylated proteins were subjected to on-bead trypsin digestion and subsequently analyzed by liquid chromatography-tandem mass spectrometry (LCMS/MS). Protein identification and quantification were carried out using Proteome Discoverer 2.4 software. Proteins that showed increased intensity in the Tim23-TurboID-V5 strain compared to the untagged control strain were considered specific. Next, protein abundance was then quantified by measuring the relative peptide intensity across three independent experiments. It is important to emphasize that many proteins were specifically labeled by the Tim23-TurboID (∼310) compared to the untagged Tim23. In this regard we identified only 37 membrane proteins, of which 18 are mitochondrial membrane proteins, as listed in Table S1. Of these we focused on Tim23 and Sfc1 related proteins. We found that Tim23 and Sfc1 were enriched proximate proteins compared to other mitochondrial control proteins such as Tim11, Tim50, Tim44 (other subunits of the TIM23 complex), Oxa1 (Mitochondrial inner membrane protein) and Aac1 (ADP/ATP Carrier (Figure 3I). These results confirm an interaction between Tim23 and Sfc1 based on TurboID.

### Proposed model

Based on these results, we propose a model for describing the interaction between Sfc1, Tim23, and succinate and their effect on mitochondrial protein import (Figure 4). Tim23 is a component of the translocase responsible for the import of matrix proteins and some inner membrane proteins. Both Tim23 and Sfc1 reside within the mitochondrial inner membrane. Our model postulates that Sfc1 and the Tim23 complex physically interact, which in turn affects the rate of protein import. In the absence of succinate, Tim23 is attached to Sfc1 resulting in slower translocation of aconitase (Figure 4A, left illustration). In the presence of succinate, Tim23 physically separates from Sfc1, facilitating protein import of aconitase into mitochondria (Figure 4B, right illustration).

**Figure 4.**
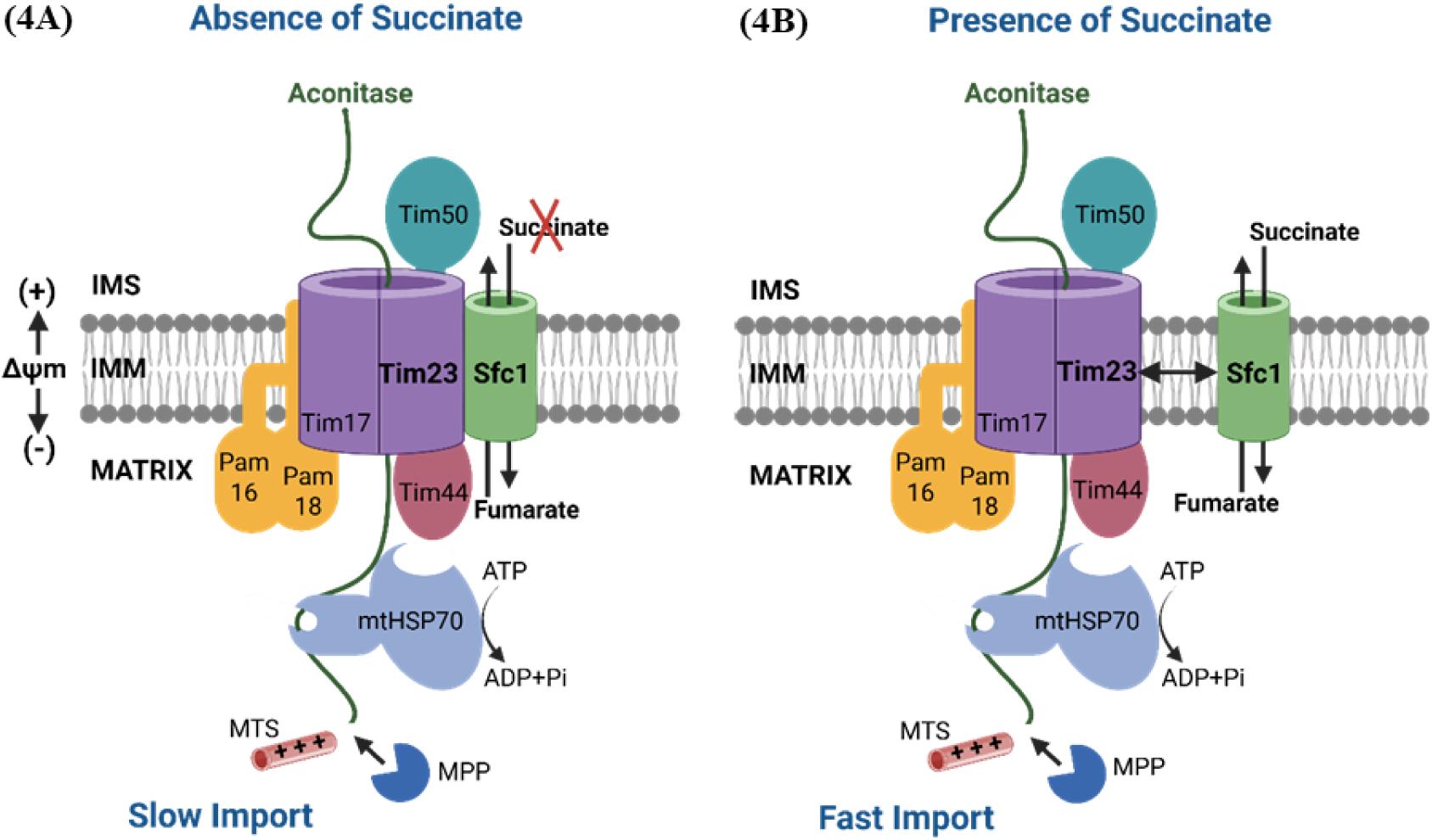
Model of the Sfc1, Tim23 and succinate interaction and the effect on mitochondrial protein import. Tim23 (a component of the translocase of the inner mitochondrial membrane) and Sfc1 (succinate-fumarate carrier) are both localized to the inner mitochondrial membrane (IMM). **(4A)** In the absence of succinate (or when Sfc1 is overexpressed), Tim23 is attached to Sfc1 resulting in a reduction in translocation of protein through Tim23. **(4B)** In the presence of succinate (or when Sfc1 is deleted), Tim23 is physically separated from Sfc1 facilitating protein import into mitochondria.

### Succinate reduces the Sfc1-Tim23 interaction

As described earlier, succinate facilitates Aco1 import into mitochondria (Figures 2A and 2B, pulse chase labeling experiments). According to the model succinate modulates protein import into the mitochondria by affecting the Sfc1-Tim23 interaction. Yeast cells expressing Sfc1 tagged with a long YFP and Tim23 tagged with a short YFP, were grown in the absence of acids (Figure 5A), the presence of 25 mM of mono-ethyl succinate (Figure 5B) or 25mM di-ethyl malate (Figure 5C) for 2hours. Indeed, we find that succinate induces the separation of Sfc1 and Tim23, resulting in a significant reduction in the percent of fluorescent cells as compared to the controls (Figure 5D). Thus, we conclude that succinate induces the dissociation of Sfc1 and Tim23, which in turn modulates import of certain proteins into mitochondria.

**Figure 5.**
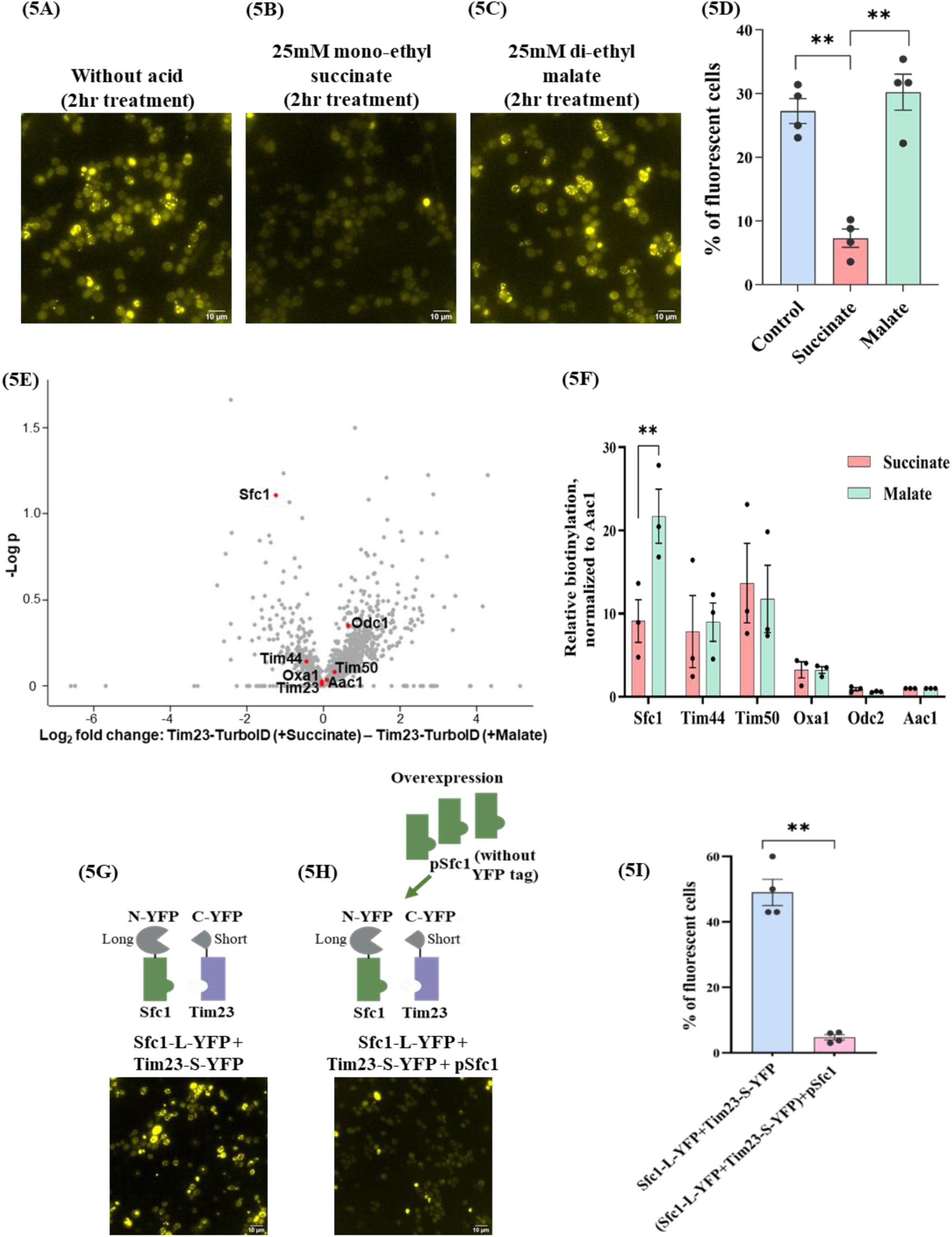
(5A) Succinate induces the dissociation of Sfc1 and Tim23. Cells expressing Sfc1-L-YFP and Tim23-S-YFP were grown in absence of acid **(5A)**, in presence of 25mM of mono-ethyl succinate **(5B)** or di-ethyl malate **(5C)** for 2 hr. According to fluorescence microscopy, succinate induces the dissociation of Sfc1 and Tim23, resulting a significant reduction in the fluorescent cells as compared to the control and malate. **(5D)** Quantification represents four independent experiments (mean ± SEM [n=4], two-tailed student’s t-test indicated statistically significant mean differences between control cells and cells treated with mono-ethyl succinate (**p=0.002) and di-ethyl malate (**p=0.006). **(5E) Succinate decreases the proximal biotinylation of Sfc1 by Tim23.** TurboID of Tim23 (fused to biotin ligase) identified modified proteins by LC-MS/MS (as described in Figure 3F). **(5E)** Volcano plot showing differential protein enrichment based on LC-MS/MS analysis. Each point represents a protein identified and quantified across conditions. The x-axis indicates the log_2_ fold change in protein intensity (Succinate vs Malate), and the y-axis shows the –log_10_ pvalue from a t-test comparing protein abundance between conditions. **(5F)** The relative levels of biotinylation for Sfc1, and other mitochondrial proteins were assessed through LC-MS/MS analysis, normalized to the control protein Aac1. Succinate significantly decreases the relative biotinylation of Sfc1 by 58% compared to malate (three independent replicate experiments, mean ± SEM [n=3], two-tailed student’s t-test **p=0.004). **(5G) Overexpressed Sfc1 competes *in vivo* with Sfc1-YFP for Tim23 binding.** Yeast cells transformed with both Sfc1 fused to long YFP and Tim23 fused to short YFP reveal significant fluorescence **(5G)**, whereas, cells co-transformed with a plasmid encoding Sfc1 (p425-Sfc1), displayed a significant reduction in the fluorescence **(5H)**. The quantification in the graph represents four independent experiments (mean ± SEM [n=4], two-tailed student’s t-test **p=0.002) **(5I)**.

To assess the impact of succinate on the Tim23-Sfc1 interaction, proximal biotinylation was employed as above. Yeast cells were cultured in biotin-depleted medium containing organic acids and then biotin was added as described in Figure 3H. Again, for the sake of brevity, the description of Tim23-TurboID-V5 expression and TurboID-based biotinylation in the presence or absence of succinate is provided in Figures S8A and S8B. Differential protein enrichment was assessed using a volcano plot based on LC-MS/MS analysis (Figure 5E). Each point represents a protein quantified across succinate and malate conditions, with the x-axis showing the log₂ fold change (succinate versus malate) and the y-axis indicating the log₁₀ p-value from a t-test. Sfc1 showed a significant decrease in biotinylation in the presence of succinate. Fascinatingly, the relative biotinylation levels of Sfc1 decreases by 58% in succinate compared to malate (Figure 5F). In contrast, control mitochondrial proteins Tim44, Tim50, Oxa1, Odc2 (Oxodicarboxylate carrier) and Aac1 did not exhibit significant changes in biotinylation, clustering near the origin of the plot. Worth mentioning is that Tim23 fused to TurboID-V5 exhibited self-biotinylation, with a lower level in the presence of succinate compared to malate, but this difference was not statistically significant (Figure S8E). We also compared samples without added acid to those with succinate (Figure S8C) and malate (Figure S8D). These control volcano plots showed insignificant changes in Sfc1 or other mitochondrial proteins, confirming that the reduction in Sfc1 biotinylation is specific for succinate. These results suggest that succinate specifically impairs the interaction between Tim23 and Sfc1, leading to decreased proximity-dependent biotinylation.

### Overexpressed Sfc1 competes for Tim23 binding

Yeast cells co-expressing Sfc1 fused to long YFP and Tim23 fused to short YFP, exhibits strong cellular fluorescence (Figure 5G). However, when these cells were co-transformed with a third plasmid encoding Sfc1 (p425-Sfc1) lacking any tags, a significant decrease in the number of fluorescent cells was observed (Figure 5H). We quantified this (Figure 5I) and found that, cells transformed with both plasmids encoding Sfc1 and Tim23 fused to YFP displayed 49% of fluorescent positive cells, while the same cells co-transformed with p425-Sfc1 without a YFP tag, exhibited a reduction in the fluorescent cells to less than 5%. This is explained by competition between the overexpressed untagged-Sfc1 and YFP-Sfc1 for Tim23 binding, which takes place *in vivo*. Again, our findings suggest that the interaction occurring within the mitochondria between Sfc1 and Tim23 is specific.

### Modification of the Sfc1 sequence affects its interaction with Tim23

At this stage of the research, we chose to examine mutations corresponding to highly biotinylated peptides of Sfc1 identified by TurboID. The notion is that these peptides within the structure of Sfc1, interact with Tim23, and their alteration may affect the interaction of these two proteins. We generated Sfc1 mutants and analyzed the effect on Sfc1-Tim23 interactions using fluorescence microscopy. In total, we created nine Sfc1 mutants and examined their expression by western blot (Figures S9A and S9B) and the amino acid residues modified in each mutant are listed in Table 1. We then examined the Sfc1-Tim23 interaction by fluorescent microscopy as described in the previous sections (Figures 6A-6J). The level of fluorescence was quantified and is presented in Figure 6K. While three of the Sfc1 display significant fluorescence six others show a significant reduction in fluorescence suggesting that in the latter the protein sequence alteration impairs the Sfc1-Tim23 interaction. Growth of yeast cells bearing Sfc1 mutants on ethanol and acetate was used as an indication of Sfc1 functionality. A yeast strain deleted for Sfc1 (Δ*sfc1*) cannot grow on ethanol and/or acetate as the sole carbon source (Figure 6L). To assess the function of Sfc1 in the above mutants, we transformed plasmids encoding wild-type Sfc1 or a Sfc1 mutant into Δ*sfc1* strains and cultured them on media with glucose, galactose, and ethanol/acetate as the sole carbon and energy source. We found that some of the Sfc1 mutants (M1-M5 and M8-M9) can replenish growth on ethanol/acetate as the sole carbon source (Figure 6L). Sfc1-mutants M7 and M6 exhibit very low fluorescence (Figures 6B and 6C) and have lost their ability to support growth on ethanol/acetate (Figure 6L and Table 1) indicating that these mutant proteins are inactive and are of less interest in this study. Sfc1-mutants M2, M4 and M5 exhibited close to wild-type fluorescence (Figures 6J, 6I and 6H, respectively) and efficient growth on ethanol/acetate (Figure 6L and Table 1) suggesting that these mutations had insignificant effects on the function of Sfc1 and on the interaction of Tim23 and Sfc1. Again, these are of less interest in this study. Exciting are Sfc1 mutants M1, M3, M8 and M9 which exhibit strong effects on fluorescence with no (M1 and M3; Figures 6E and 6G) or partial (M8 and M9; Figures 6F and 6D) effects on the growth on ethanol/acetate (Figure 6L and Table 1). Worth mentioning is that while Sfc1 mutants M8 and M9 harbor deletions of amino acid codons, Sfc1 mutants M1 and M3 contain amino acid substitutions. Thus, this latter group of mutants indicate that we can affect the interaction of Sfc1-Tim23 interaction without losing the transporter’s activity (succinate-fumarate exchange).

**Table 1:**
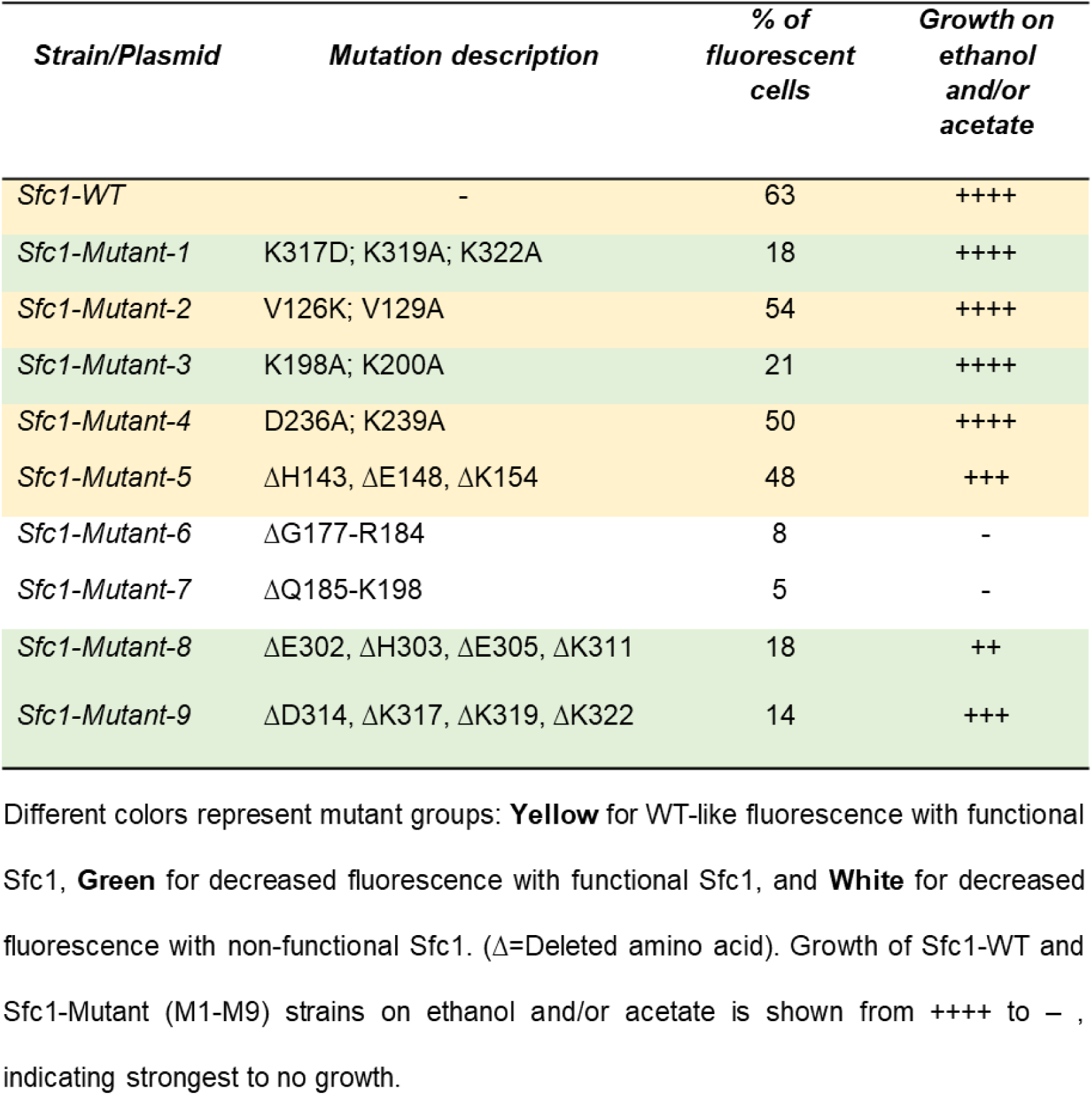
Sfc1 mutants employed in the study:

**Figure 6.**
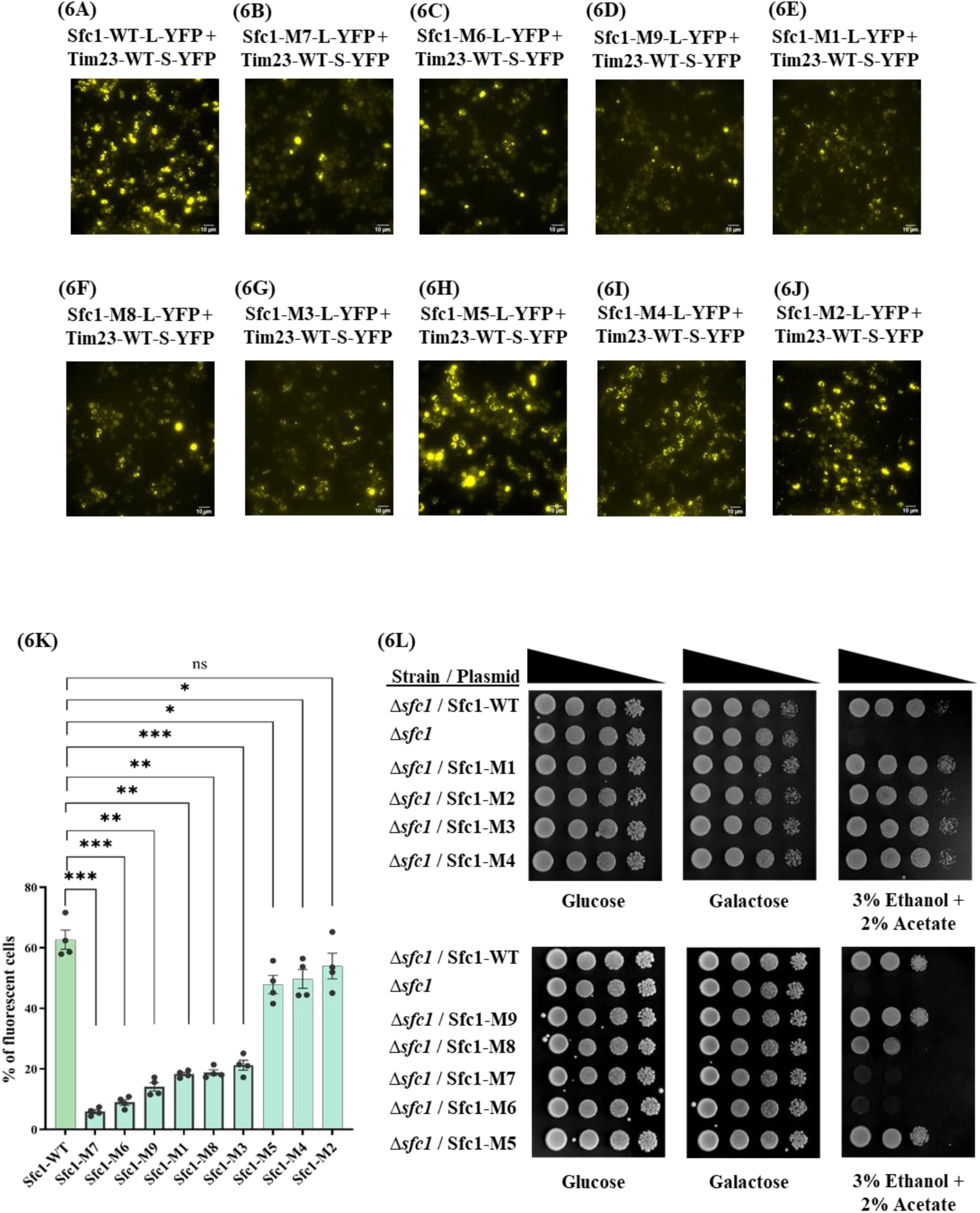
Modification of the Sfc1 sequence affects the Sfc1-Tim23 interaction. Yeast cells transformed with both WT-Sfc1 fused to long YFP and WT -Tim23 fused to short YFP **(6A),** exhibited significant fluorescence (Sfc1-WT-L-YFP + Tim23-WT-S-YFP). The Sfc1 mutants (Sfc1-Mutant-X-L-YFP + Tim23-WT-S-YFP) are referred to by number as follows and details can be found in Table 1: Sfc1-M7 **(6B)**, Sfc1-M6 **(6C)**, Sfc1-M9 **(6D)**, Sfc1-M1 **(6E)**, Sfc1-M8 **(6F)**, Sfc1-M3 **(6G),** Sfc1-M5 **(6H)**, Sfc1-M4 **(6I)**, Sfc1-M2 **(6J)** fused to long YFP and WT-Tim23 fused to short YFP (M: mutant number). The quantification in the graph **(6K)** represents the percentage of fluorescent cells out of the total cell count, based on data from four independent experiments (mean ± SEM [n=4], two-tailed t-test *p<0.05, **p<0.01, and ***p<0.001). **(6L) Certain Sfc1 mutants replenish growth on ethanol and acetate medium.** WT-Sfc1 and Sfc1-mutant strains (M: mutant number) were transformed into Sfc1 knockout strains (Δ*sfc1*) and grown on glucose, galactose, and ethanol-acetate medium plates to assess the restoration of growth. While Δ*sfc1* strains are unable to grow with ethanol-acetate as the sole carbon source, most Sfc1 mutants still exhibited robust growth under these conditions (except Sfc1-M6 and Sfc1-M7) indicating that most Sfc1 mutants retain a functional Sfc1.

### Modification of the Tim23 sequence affects its interaction with Sfc1

In a parallel approach, we attempted to identify residues of Tim23 crucial for its interaction with Sfc1, by site-directed mutagenesis guided by the TurboID analysis of Tim23. We introduced mutations within the Tim23 sequence and then analyzed Tim23-Sfc1 interaction by fluorescence microscopy. We generated a total of four Tim23 mutants in which we created deletions of four to five amino acids. Three out of four of these Tim23 deletion mutants affected the Tim23-Sfc1 interaction, resulting in a significant decrease in the number of fluorescent cells (Figure S10). We verified the expression of both wild-type Tim23 and Tim23 mutants (M1-M4) fused to the short YFP fragment by western blot analysis using an anti-Tim23 antibody (Figure S9C, lower panel) and an Aco1 antibody as an internal control (top panel). As a control, yeast cells transformed with two plasmids, one encoding WT-Tim23 fused to short YFP and the other encoding WT-Sfc1 fused to long YFP, displayed significant fluorescence (Figure S10A). Cells transformed with plasmids encoding either Tim23-M1, or Tim23-M2, or Tim23-M4 fused to short YFP, along with WT-Sfc1 fused to long YFP, exhibited a highly significant reduction in fluorescence (Figures S10B, S10C and S10E, respectively); 4%, 7%, and 9% fluorescent-positive cells, respectively versus 56% of the Tim23 wild-type (Figure S10A). The fluorescence reduction in Tim23-M3 was not statistically significant and closely resembled that of WT-Tim23 (Figure S10D). The specific amino acid residues altered in each mutant are listed in Table S2. These findings suggest that the alterations in the Tim23 sequences of mutants M1, M2, and M4 impact the Tim23-Sfc1 interaction, however, in contrast to the previous section, here we do not know whether the Tim23 mutants are active in protein translocation.

### Rosetta-MP Docking of Sfc1 and Tim23

As discussed above the mutagenesis of Sfc1 and Tim23 was based on peptides of Sfc1 and Tim23 identified by TurboID. The fact that such mutations affect the interaction of Sfc1 and Tim23 supports our model of their interaction. Nevertheless, the precise interface and the location of mutations which reduce binding of the two complexes, is unknown. To create a platform for thought, we employed an implicit membrane-based docking method to determine the residues that are involved in binding at the interface. With the sequence of Sfc1 and Tim23 as the input, we have used AlphaFold3 (AF3) ^33^ to predict the structure of the dimer, and then refined the top-ranking models using the Rosetta based flexible membrane protein docking method ^34^. Our assumption that the interface of the complex lies in the transmembrane region, is based on an AF3 predicted structure of the Sfc1-Tim23 complex, which was subsequently positioned into the membrane using the PPM webserver ^35^.

For this study, we have employed our recently developed Rosetta-MPDock, a flexible transmembrane (TM) protein docking protocol designed to capture binding-induced conformational changes ^34^. In addition, to using ensembles generated by Rosetta-based methods such as relax, back-rub and normal mode analysis (NMA), we incorporated DL-based methods to generate ensembles of monomer backbones ^36^, which provided higher backbone sampling. The top 5 structures predicted by AF3 were used as input poses for Rosetta-MP docking, to generate 8000 docked decoys for each. We subsequently ranked these decoys based on their low interface and total scores, as calculated using membrane informed energy function ^37–39^ and selected the top 20 decoys for each input pose. Through clustering of these top scoring docked structures, we identified a distinct interface shown in Figure 7A. A detailed analysis of the interface was conducted to determine the nature of interactions between the two chains. Figure S11 shows the distance map overlaid with the residues identified by TurboID and mutagenesis. The residues involved in the interaction between two chains, along with the type of interactions are summarized in Table S3. The interactions are primarily due to hydrophobic and electrostatic interactions.

**Figure 7.**
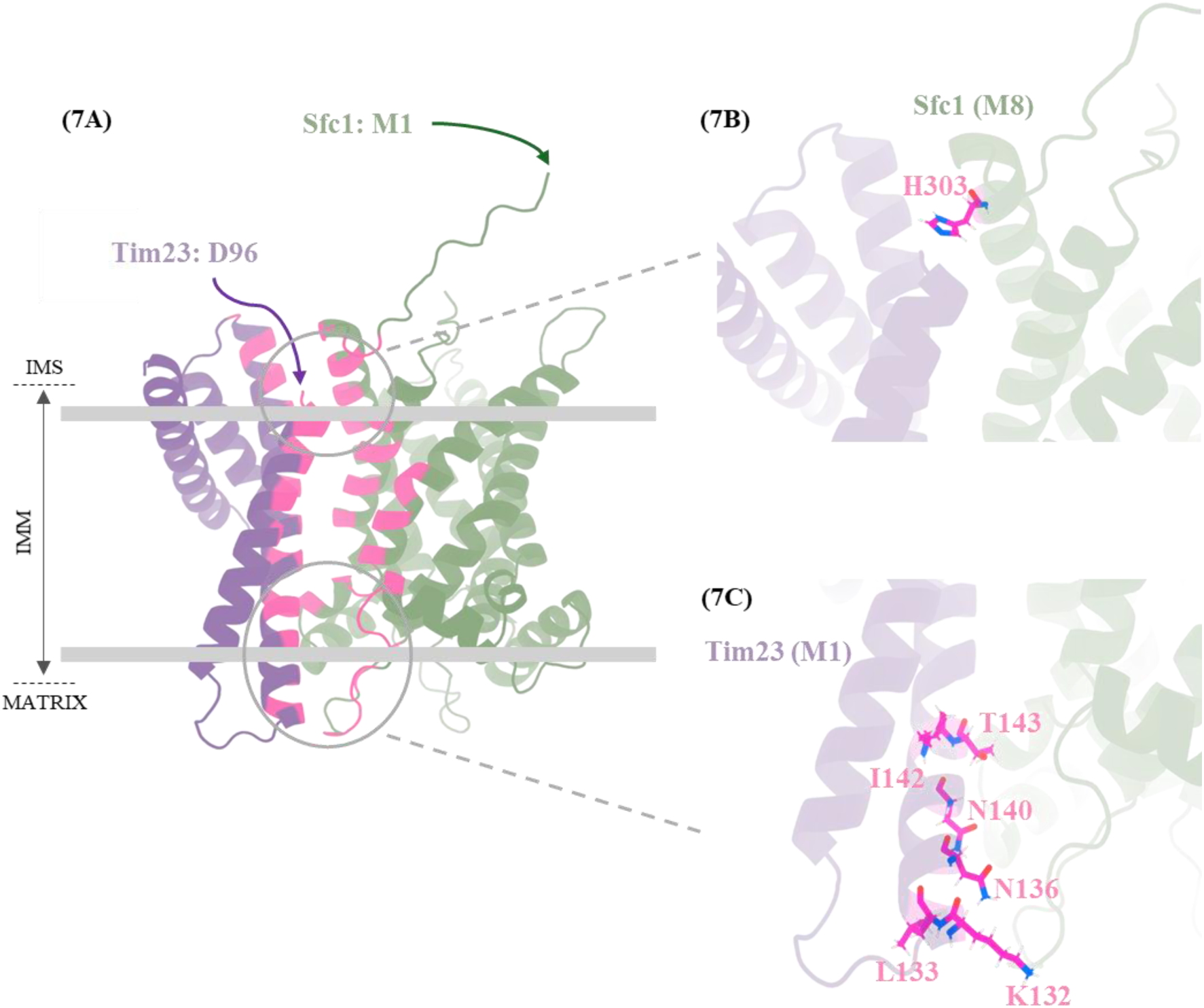
Structural insights into the Sfc1 -Tim23 complex using RosettaMPDock, an implicit membrane-based flexible protein docking method. **(7A)** A distinct low-energy interface was identified through extensive conformational sampling using RosettaMPDock, followed by ranking based on membrane-informed energy functions. The structures of Tim23 (shown in purple) and Sfc1 (in green) were predicted using Alphafold3 (AF3) and subsequently refined through Rosetta-based flexible membrane protein docking protocol. Interface residues are highlighted here in pink. **(7B)** Residues in the experimental Sfc1 mutant M8 that are located at the Sfc1-Tim23 interface. **(7C)** Interface residues of the Sfc1-Tim23 complex corresponding to the experimental Tim23 mutant M1.

Revisiting the TurboID-based mutagenesis, we overlaid the mutated residues onto the distance map of the interface (as shown in Figures 7B-7C, Rosetta-MP Dock predicted interface aligns with residues in a Sfc1 mutant (Sfc1-M8) and residues in a Tim23 mutant (Tim23-M1). The significant decrease in fluorescence in the mutants such as Sfc1 mutant M8 (Figure 6F) and Tim23 mutant M1 (Figure S10B) verifies a perturbation in the interaction sites of Sfc1 and Tim23. These results show that Sfc1 mutant M8 (Figure 7B) and Tim23 mutant M1 (Figure 7C) are positioned at the interface of the Sfc1-Tim23 model.

## Discussion

Metabolite signaling is an important aspect of cellular function and regulation, however, we do not always understand how this is achieved. Here we show for the first time how the flow of a metabolite through a carrier in a membrane, affects the interaction of this carrier with a different membrane complex, thereby affecting the latter’s activity. Looking back, previous studies in our lab discovered that addition of succinate (a product of the glyoxylate shunt) or elimination of the glyoxylate shunt in yeast, alters the distribution of fumarase and aconitase, increasing the levels of the mitochondrial versus the cytosolic isoforms ^9^. It is crucial to understand that succinate does not affect the subcellular distribution of all mitochondrial proteins but rather certain proteins, including aconitase and fumarase. Our group discovered that both tricarboxylic acid (TCA) cycle enzymes fumarase (Fum1) and aconitase (Aco1) in the yeast *Saccharomyces cerevisiae* are dual targeted to the cytosol and mitochondria by a reverse translocation mechanism ^1–8^. Fumarase folding drives its retrograde return to the cytosol, with rapid folding blocking efficient post-translational import into mitochondria in both *in vivo* and *in vitro* analyses ^5,8^. Moreover, mutations that alter fumarase conformation leave mitochondrial targeting intact but block its reverse translocation, while overexpression of a cytosolic Hsp70 shifts fumarase toward mitochondria and reduces mitochondrial Hsp70 impairs full import and increases its cytosolic pool ^6^. Accordingly, mass spectrometry showed that cytosolic and mitochondrial fumarase are identical and lack post translational modifications, supporting the conclusion that folding alone drives its distribution ^6^. In the case of aconitase, N-C terminal domain interaction was crucial for efficient post-translational import and dual targeting, with the C-terminal domain providing a chaperone like function that was modulated by cytosolic Hsp70 (Ssa1/2) ^4,12^. Thus, these proteins are special in that they have been shown to be able to move forwards and backwards in the translocation apparatus and this results in their dual targeting into and out of mitochondria. The precise mechanism of how Sfc1 and succinate affect translocation and reverse translocation of fumarase and aconitase remains to be fully elucidated. The connection between these stories is that Sfc1, which is a mitochondrial succinate-fumarate carrier, is also required for the glyoxylate shunt. Sfc1 mediates the transport of succinate from the cytosol into the mitochondria, coupled with the export of fumarate from the mitochondria to the cytosol. The exported fumarate is subsequently directed into the gluconeogenic pathway, and Sfc1 is essential for the utilization of ethanol/acetate as a carbon source ^16,21,22^. Essentially, knockout of the *SFC1* gene (Δ*sfc1*), changes the subcellular distribution of fumarase resulting in lower levels of fumarase in the cytosol and higher levels in the mitochondria. Accordingly, Sfc1 is also crucial for regulating the subcellular distribution of aconitase, as yeast lacking the *SFC1* gene exclusively localize aconitase to mitochondria.

The objective of this study was to understand how Sfc1 and succinate affect mitochondrial protein import and dual targeting in yeast. The results obtained from pulse-chase experiments indicate that succinate and Δ*sfc1* facilitate aconitase (Aco1) import into mitochondria, while overexpression of Sfc1 slows down Aco1 import. The model presented here (Figure 4) is that Sfc1 and the Tim23 complex (the protein translocase of the mitochondrial inner membrane) can physically interact, which in turn regulates protein import. The model predicts that in the absence of succinate or when Sfc1 is over-expressed, Sfc1 is attached to Tim23, leading to a slower rate of protein translocation. However, in the presence of high concentrations of succinate or a Sfc1 knockout (Δ*sfc1*), Sfc1 and Tim23 dissociate resulting in higher protein import. To test this hypothesis, protein-protein interactions between Sfc1 and Tim23 were analyzed by Co-immunoprecipitation (Co-IP), Bimolecular Fluorescence Complementation (BiFC) techniques and Biotin-Based Proximity labeling (TurboID). The results revealed direct interaction between Sfc1 and Tim23, and more importantly, the specificity and non-random nature of the interaction between these two, was confirmed. Furthermore, addition of succinate induces the dissociation between Tim23 and Sfc1, which can modulate import of certain proteins into mitochondria as evaluated using fluorescence microscopy and TurboID. In the next step, we evaluated the Sfc1-Tim23 interaction through mutagenesis of potential binding sites, guided by the TurboID results, and visualized the changes using fluorescence microscopy. Mutations in Sfc1 or Tim23 disrupted their interaction, as shown by a reduced number of fluorescent cells, thereby identifying potential interaction sites. Using these insights, we generated a model of the Sfc1-Tim23 complex with Rosetta-MP Docking ^34^, based on the TurboID and mutagenesis data. These observations imply that both Sfc1 and succinate, regulate yeast mitochondrial protein import and dual targeting, which is crucial to balance cellular processes such as glyoxylate shunt, TCA cycle, gluconeogenesis etc.

The interaction of entities in the membrane which are responsible for transfer of molecules through or into the membrane is a new and important issue. Membrane protein-protein interactions have been reported in the past and this is also true for the TIM23 complex which has been shown to interact with the bc1 (cytochrome *bc*1) and the COX (cytochrome *c* oxidase) complex ^40–42^. Interestingly, Pet9 (Aac2) an ADP/ATP carrier isoform, exists in physical association with the cytochrome *bc*1-COX complexes and the TIM23 machinery and presumably brings about the association of these complexes ^43^. Other studies indicate protein-protein interactions within membranes, however, the functional relevance of these interactions remains to be elucidated ^44–47^. Tim23 has long been considered as the channel-forming subunit, with Tim17 providing structural support ^48–50^, but recent findings challenge this model ^51,52^. Structural and biochemical studies revealed a dynamic Tim23-Tim17-Mgr2 heterotrimer, and cryo-EM data shown that their soluble domains are flexible ^52^. Tim23 also undergoes voltage-dependent conformational changes via its aqueous-facing TM2 helix ^11,53,54^. However, it remains debated whether Tim23 or Tim17 individually or together form the protein conducting channel, or whether multiple copies of these subunits contribute at different stages of protein translocation ^52^. As pointed out at the beginning of this discussion, here we show how the flow of a metabolite through a carrier in a membrane, affects the interaction of this carrier with a different membrane complex, thereby affecting the latter’s activity. This is a novel perspective for metabolic signaling in the cell. In this regard when we open books on biochemistry and molecular biology, we find that metabolites are defined as small molecules that are intermediates and building blocks of cells. These molecules can regulate enzyme activity, participate in the regulation of gene expression and the generation of energy. Here we show a new feature of metabolic signaling; the flow of a metabolite, in this case succinate, through its membrane carrier, Sfc1, impacts the translocation of a protein through the Sfc1-attached TIM23 complex.

## Materials and Methods

### Yeast strains and growth conditions

*Saccharomyces cerevisiae* yeast strains were obtained from Euroscarf; BY4742 (MATα his3Δ1 leu2Δ0 lys2Δ0 ura3Δ0), Δ*sfc1* (BY4742; Mat α: his3Δ1; leu2Δ0; lys2Δ0; ura2Δ0) and pSfc1 (pSfc1 overexpression plasmid under GAL promoter transformed into BY4742). Strains were cultured at 30°C or as specified in synthetic depleted (SD) medium containing 0.67% (w/v) yeast nitrogen base without amino acids, 2% glucose/ 2% galactose (w/v)/ 2% Acetate (w/v)/ 3% Ethanol (v/v), supplemented with the necessary amino acids (50 µg/ml). For solid medium, 2% agar was added. X-gal plates were prepared by adding a stock solution containing 2% galactose, 1% raffinose, 0.008% X-gal (dissolved in 100% N, N-dimethylformamide), and 1x BU salts (25 mM sodium phosphate buffer, pH 7.0) to an autoclaved medium.

### Plasmids and primers

Plasmids used in this study are listed in Table S4. Constructs were generated using restriction cloning or Gibson assembly and verified by Sanger sequencing. Mutants were generated via site directed mutagenesis (Bio Basic Asia Pacific). Primers used for cloning are listed in Table S5.

### α-Complementation assay

Yeast cultures containing plasmids encoding the specified α fusion proteins, along with either pωc or pωm, were generated^27,55^. The resulting colonies were grown on plates containing 80 mg/ml X gal and incubated in the dark at 30°C for 72 hours.

### Subcellular fractionation

Yeast cultures were grown to an OD600 of 1.5, and mitochondria were isolated as previously described ^8,56^. Spheroplasts were generated using Zymolyase-20T. To assess cross-contamination in our subcellular fractionation experiments, we used anti-Hsp60 as a mitochondrial marker and anti-hexokinase1 (anti-HK) as a cytosolic marker. The intensity of the cytosolic and mitochondrial bands was quantified densitometrically using ImageJ software.

### Pulse-chase and immunoprecipitation

Yeast cultures were grown in 25 ml of galactose medium containing 1/20 of the total volume of 1M K-Phosphate buffer (pH 6.8) at 30°C until reaching an OD600 of 1.2. The cells were then collected by centrifugation and resuspended in 25 ml of fresh Gal-Met medium with K-Phosphate buffer, followed by incubation at 30°C for 1 hour to deplete methionine. Methionine-depleted cells were treated with 0.03 mM CCCP for 1.5 minutes and labeled with 80 µCi of [35S] methionine, then incubated at 30°C for 15 minutes. Afterward, the cells were treated with 0.1 M DTT, and labeling and translation were stopped by adding excess cold 0.003% methionine, 0.004% cysteine, and 0.001% cycloheximide. The cells were further treated with 10 mM mono-ethyl succinate and di ethyl malate in the presence of 1M K-Phosphate buffer and incubated at 30°C for 0, 5, 10, and 15 minutes. At the indicated times, 400 µl samples were taken, placed on ice, and treated with 10 mM sodium azide for washing. Labeled cells were collected by centrifugation, resuspended in 10 mM Tris/EDTA buffer (pH 8.0) containing 1 mM PMSF, and disrupted using glass beads by shaking for 20 minutes at 4°C. The resulting supernatant fraction was obtained by centrifugation, denatured by boiling in 1% SDS, and subjected to immunoprecipitation using anti-aconitase rabbit antiserum and magnetic Dynabeads Protein A (Invitrogen) in TENN buffer (200 mM Tris, pH 8.0; 20 mM EDTA, 2% NP-40, 600 mM NaCl). The samples were analyzed by SDS-PAGE, followed by phosphor imaging and autoradiography for visualization. Band intensities of the precursor (p) and mature (m) forms were quantified by densitometry using TINA 2.09 software.

### Protein import and detection in isolated mitochondria

Mitochondria were isolated from yeast following the method outlined by Daum et al. 1982 ^56^. Protein synthesis was carried out using [35S] methionine in rabbit reticulocyte lysate (Promega). The import of aconitase (Aco1) into isolated mitochondria was carried out using a HEPES-containing buffer (0.1 M HEPES, pH 7.3; 1.2 M Sorbitol, 0.16 M KCl, 10 mM MgCl_2_, 0.25 M Methionine, 100 mM ATP, and 100 mM NADH). A 2.2 mg aliquot of isolated mitochondria was resuspended in 200 µl of import buffer. The suspension was divided into three equal parts and incubated with either 10 mM potassium succinate, 10 mM potassium malate, or 10 mM potassium phosphate buffer (KPB), pH 7. S³⁵-radiolabeled Aco1 was produced via *in vitro* transcription/translation using reticulocyte lysate (Promega). The import reaction was initiated by adding S³⁵-labeled Aco1 to the import buffer containing the isolated mitochondria. Following incubation at 25°C for 0 to 16 minutes, the samples were split into two 100 µl aliquots. One set was treated with 1 mg/ml proteinase K for 10 minutes on ice, and the reaction was stopped by adding 2 mM PMSF and 20 µM CCCP. Mitochondrial proteins were then separated by SDS-PAGE, and the S³⁵-labeled Aco1 was detected using a phosphor imager and autoradiography.

### Co-Immunoprecipitation (Co-IP)

Yeast cultures were grown in SD galactose medium to an A600 of 1.5, harvested, washed, and lysed in CHAPS buffer (50 mM Tris-Cl pH 7.5; 150 mM NaCl, 1% CHAPS, 1 mM PMSF) using glass beads. Lysates were clarified by centrifugation (5500 g, 4 °C). Protein A Dynabeads (Invitrogen) were incubated with anti-Sfc1 antisera (10 μl/50 μl beads) in PBST for 30 min at room temperature, then mixed with the clarified lysate (100–1000 μl) for 1 h. After magnetic separation, the supernatant (FT) was collected, and bound proteins were eluted with lysis buffer and 4× sample buffer (E). Immunoprecipitation with anti-Sfc1 and co-IP of Tim23 were analyzed by SDS-PAGE and western blot using anti-Tim23 antibodies.

### Fluorescence microscopy and mitochondrial labeling

Plasmid combinations listed in Table S6 were co-transformed into *S. cerevisiae* BY4742 using the Frozen-EZ Yeast Transformation II Kit for fluorescence microscopy. Transformed yeast cells were grown in appropriate filtered medium at 30 °C with shaking until they reached an OD₆₀₀ of 1.0-1.2.

For MitoTracker staining, cells were harvested by centrifugation (4500 g, 5 min) and resuspended in prewarmed medium containing 200 nM MitoTracker Red CMXRos (Invitrogen). After a 15 min incubation at room temperature, cells were washed three times with PBS, centrifuged, and concentrated tenfold. An aliquot was mounted on a glass slide, covered with a coverslip, and visualized using a 100× Olympus IX83 fluorescence microscope. Images were processed using OLYMPUS OlyVIA software.

### TurboID-based proximity labeling for pulldown of biotinylated proteins

Based on Branon et al. (2018)^32^, yeast cells were transformed using the Frozen-EZ Yeast Transformation II Kit (Zymo Research) and *LEU2*-positive transformants were selected and cultured in SD-Leu medium at 30 °C. Protein expression was induced by diluting saturated cultures (1:100–1000) into 10% SD/G-Leu (SD-Leu medium with 90% of dextrose replaced with galactose) or biotin-depleted medium (1.7 g/L YNB-Biotin-Amino acid, 5 g/L ammonium sulfate, 2 g/L dextrose, 18 g/L galactose, complete amino acids, 790 ng/mL d-biotin), followed by 18–24 h incubation at 30 °C. For Western blot, wild-type and Tim23-TurboID-V5 cells were grown in biotin-depleted medium for 16 h, then supplemented with 50 µM biotin, 1 mM ATP, and 5 mM MgCl₂ for 2 h. For organic acid treatments, 50 mM succinic or malic acid in 1 M KPB (pH 7) was added with biotin. ‘Omit biotin’ controls were grown without biotin overnight. Cells were washed five times with cold PBS and centrifuged at 4500×g for 5 min at 4 °C. Proteins were extracted using glass beads in 1% CHAPS buffer at 4 °C, then centrifuged at 5500×g for 5 min. Pellets were resuspended in CHAPS buffer and SBx4, heated at 95 °C for 5 min, and separated by 12% SDS-PAGE. For biotinylated protein enrichment, 100 µL streptavidin beads (Dynabeads M280, Invitrogen) were washed and incubated with 50 µg lysates for 1 h at room temperature, then rotated overnight at 4 °C. Beads were washed with PBS, and 5% of beads were eluted in SBx3 buffer containing 20 mM DTT and 2 mM biotin. Eluted proteins were confirmed by SDS-PAGE before LC-MS/MS.

### Trypsin digestion of biotinylated proteins on beads

The proteins attached to the beads were reduced with 10mM DTT in 8M Urea, 100mM ABC (60°C for 30 min), modified with 35mM iodoacetamide in 100mM ammonium bicarbonate (in the dark, room temperature for 30 min) and digested in 2M Urea, 25mM ammonium bicarbonate with modified trypsin (Promega) at a 1:100 enzyme-to-substrate ratio, overnight at 37°C.

### Liquid chromatography and mass spectrometry (LC-MS/MS)

The tryptic peptides were removed from the beads, desalted using homemade C18 stage-tips, dried and re-suspended in 0.1% Formic acid. The resulted peptides were analyzed by LC-MS/MS using a Q- Exactive plus mass spectrometer (Thermo) fitted with a capillary HPLC (easy nLC 1000, Thermo-Fisher). The peptides were loaded onto a C18 trap column (0.3 x 5mm, LC-Packings) connected on-line to a homemade capillary column (20 cm, 75-micron ID) packed with Reprosil C18-Aqua (Dr. Maisch GmbH, Germany) in solvent A (0.1% formic acid in water). The peptides mixture was resolved with a (5 to 28%) linear gradient of solvent B (95% acetonitrile with 0.1% formic acid) for 60 minutes followed by gradient of 15 minutes of 28 to 95% and 15 minutes at 95% acetonitrile with 0.1% formic acid in water at flow rates of 0.15 μl/min. Mass spectrometry was performed in a positive mode using repetitively full MS scan followed by collision induces dissociation (HCD) of the 10 most dominant ions (>1 charges) selected from the first full MS scan. The mass spectrometry data was analyzed using Proteome Discoverer 2.4 software with Sequest (Thermo) search engine against the *Saccharomyces cerevisiae* proteome from the Uniprot database with mass tolerance of 20 ppm for the precursor masses and 0.05 amu for the fragment ions. Oxidation on Met was accepted as variable modifications, carbamidomethyl on Cys was accepted as static modifications. Minimal peptide length was set to six amino acids and a maximum of two miscleavages was allowed. Protein-level false discovery rates (FDRs) were filtered to 1% using the target-decoy strategy. Protein table was filtered to eliminate the identifications from common contaminants and single -peptide identifications. Semi quantitation was done by calculating the peak area of each peptide based its extracted ion currents (XICs) and the area of the protein is the average of the sum of all the identified peptides from each protein. Alternatively, the data was analyzed using the Maxquant 2.1 using similar parameters.

### Rosetta-MPDock

In this study, we have utilized our recently developed Rosetta-MPDock, a flexible transmembrane (TM) protein docking protocol designed to capture binding-induced conformational changes ^34^. Rosetta-MPDock samples large conformational ensembles of flexible monomers and docks them within an implicit membrane environment. In a local docking scenario - where membrane protein partners are positioned ≈10 Å apart in the membrane in their unbound conformations, Rosetta MPDock successfully predicts the correct interface (success defined as achieving 3 near-native structures in the 5 top-ranked models) for 67% moderately flexible targets and 60% of the highly flexible targets, a substantial improvement from the existing membrane protein docking methods. Further, by integrating AlphaFold2-multimer for structure prediction with Rosetta-MPDock for docking and refinement, we achieved an improved success rate over the benchmark targets from 64% to 73%. For this work, we have used AF3 ^33^ predicted models for the complex as the starting structures, followed by Rosetta-MPDock to dock, refine and rank the structures with membrane informed score functions ^33,37–39^. In addition to using ensembles generated by Rosetta-based methods such as relax, back-rub and normal mode analysis (NMA), we incorporated ensembles of monomer backbones produced by DL based ensemble generators ^57^, which provided higher backbone sampling. For each starting structure of the complex predicted by AF3, we have generated 8000 docked decoys using Rosetta-MPDock. From these decoys, we selected the top 20 decoys for each starting structure based on their low interface score and total scores. Additional analysis has been performed on clusters of similar interfaces. Previously, the clusters were selected based on visual assessment, but we are now employing distance-based clustering techniques to make the process more systematic and rational.

## Statistical Analysis

Statistical analysis was performed using a two-tailed paired Student’s *t*-test. Data are presented as the mean ± SE from three independent experiments. Graphs were generated with GraphPad Prism 10.4.1.

## Data availability

The data supporting the findings of this work are available within the paper and/ or its supplementary information or on request from the corresponding author [O.P].

## Acknowledgements

This work was supported by grants of J.H and O.P. from the German-Israeli Foundation for Scientific Research and Development (GIF; Grant No. 1561) and the German-Israeli Project Cooperation (DIP; Grant No. 17516). R.S. received financial support from a start-up fund provided by the University of South Florida. Computational resources were provided by the CIRCE HighPerformance Computing Cluster. The funding agencies had no role in the study design, data collection, data analysis, or the decision to publish this work. We thank Prof. Doron Rapaport (Interfaculty Institute of Biochemistry, University of Tübingen) for valuable discussions and for generously providing plasmids containing split YFP tags. The illustrations were created using BioRender (https://app.biorender.com/).

## Author Contributions

K.D.: conceptualization, investigation, methodology, validation, visualization, formal analysis, data curation, project administration, writing-original draft, writing review and editing. R.S.: methodology, formal analysis, visualization, resources, writing-review and editing. T.Z.: visualization, formal analysis, writing-review and editing. J.M.H.: conceptualization, writing-review and editing. B.K.: conceptualization, investigation, project administration, supervision. O.P.: conceptualization, resources, project administration, supervision, writing-original draft, writing-review and editing, funding acquisition.

## Supplemental Information

**Figure S1.**
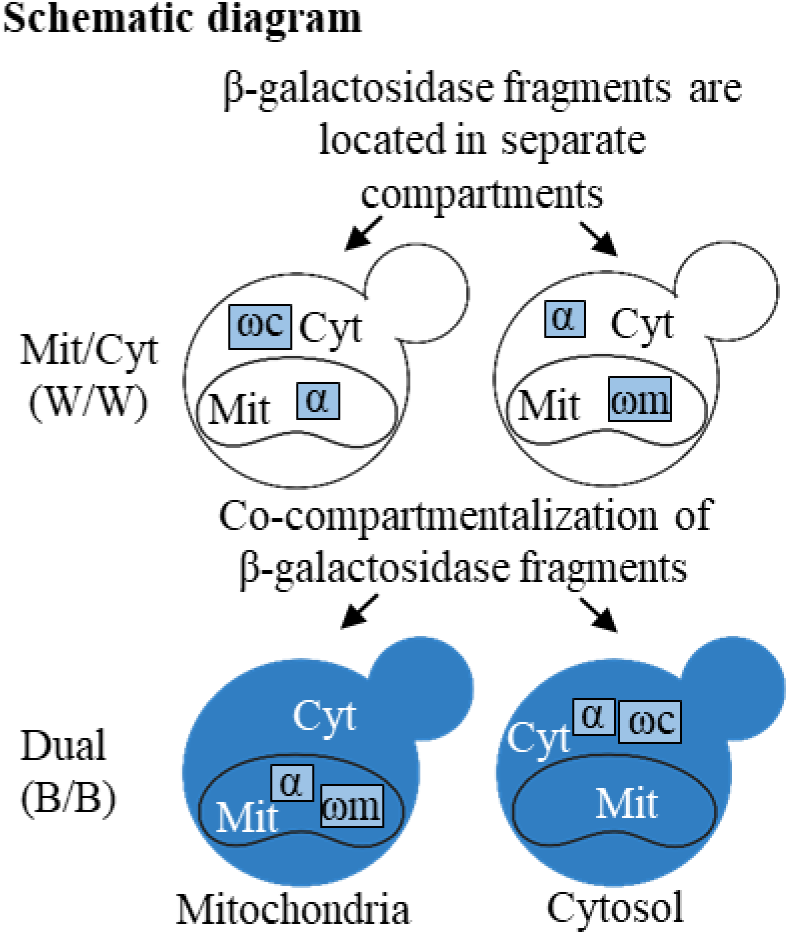
Schematic diagram of the α-complementation assay: Co-localization of the βgalactosidase fragments, α and ω, within the same compartment (bottom panel) allows assembly of the active enzyme and results in blue colonies on X-gal plates. Conversely, if the two fragments are in separate compartments, this will result in white colonies (top panel). Cyt, cytosol; Mit, mitochondria; W, white; B, blue.

**Figure S2.**
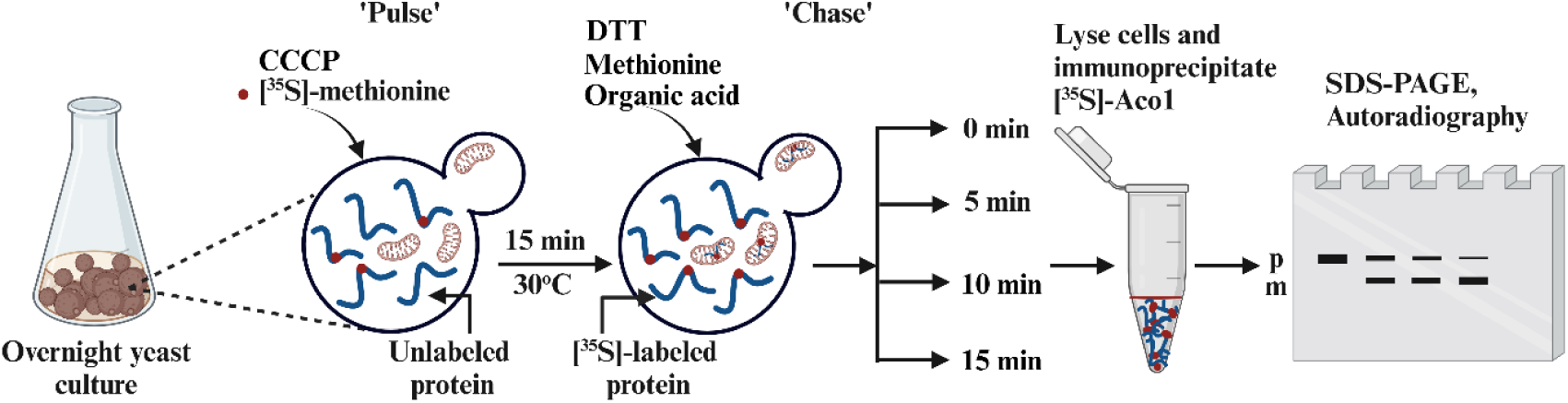
Illustration depicting the *in vivo* pulse-chase assay. Wild-type yeast cultures, induced in galactose medium, were labeled for 15 minutes with [^35^S]- methionine in the presence of CCCP to accumulate labeled precursors. The labeled cultures were then chased with DTT, an excess of cold methionine in the absence of acid, or in the presence of 10 mM mono-ethyl succinate or in the presence of di-ethyl malate at 30°C for 0, 5, 10, and 15 minutes. Samples were collected at the specified time points, immunoprecipitated with an anti-Aco1 antibody and analyzed by SDS-PAGE followed by autoradiography.

**Figure S3.**
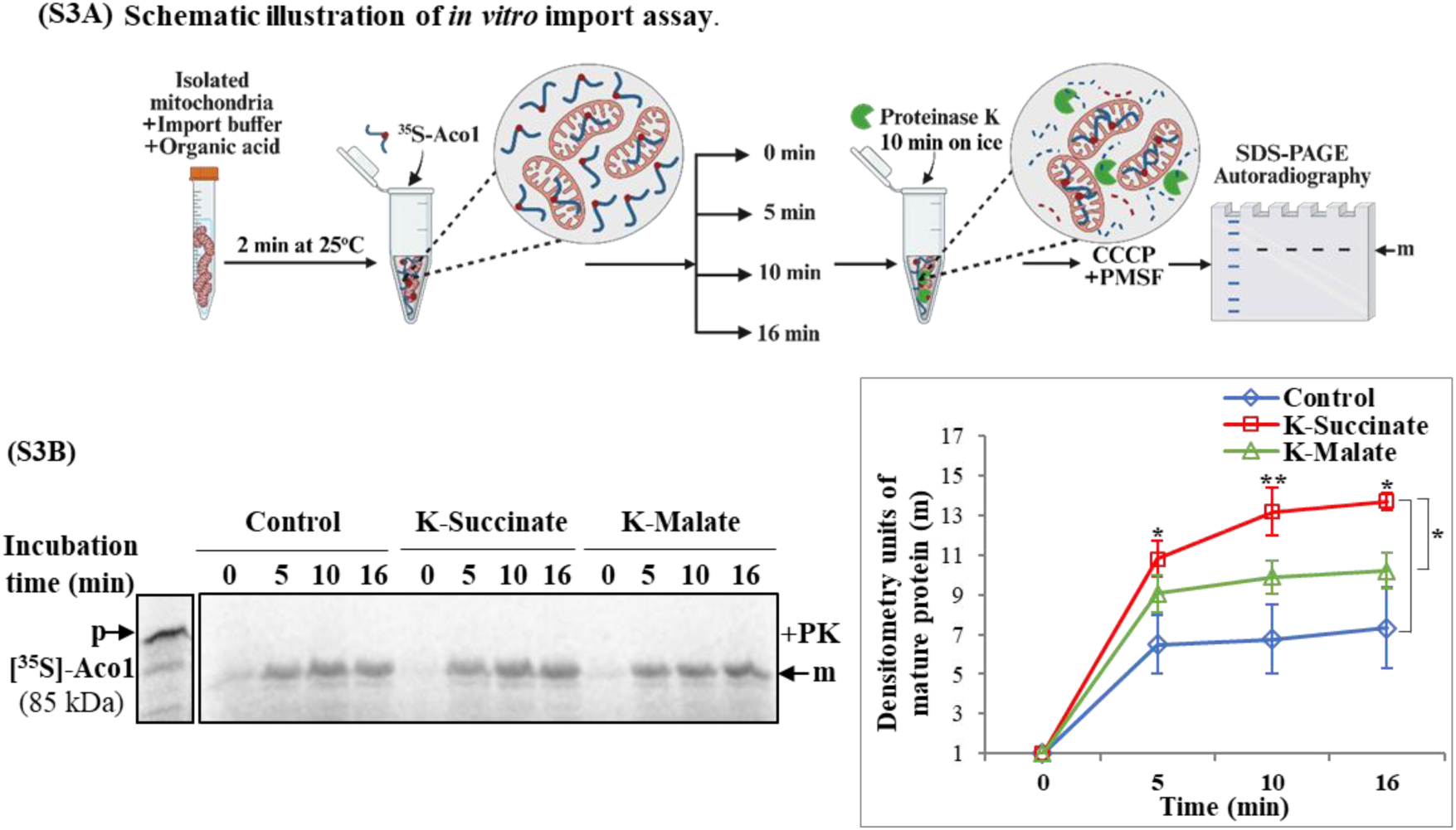
(S3A) Schematic illustration of *in vitro* import assay. [^35^S]-methionine labeled Aco1 was transcribed and translated *in vitro* in reticulocyte lysate. The import process commenced by adding [^35^S]-methionine labeled Aco1 to an import buffer containing isolated mitochondria and in the presence or absence of organic acid. At specified intervals, samples were withdrawn and subsequently incubated with proteinase K to digest proteins located on the mitochondrial surface. The [^35^S]-labeled Aco1 was then analyzed by SDS-PAGE and autoradiography as a function of time. **(S3B) The effect of succinate and malate on Aco1 import into isolated mitochondria *in vitro*.** The [^35^S] autoradiography on the left panel shows the processed Aco1 (m−mature) inside mitochondria, after treatment of the mitochondria with proteinase K under various conditions and right panel shows quantification. The amount of processed Aco1 (m−mature) is plotted against time. The quantification shown in the graph represents data from three independent experiments (mean ± SEM [n=3], twotailed t-test *p < 0.05, **p < 0.01).

**Figure S4.**
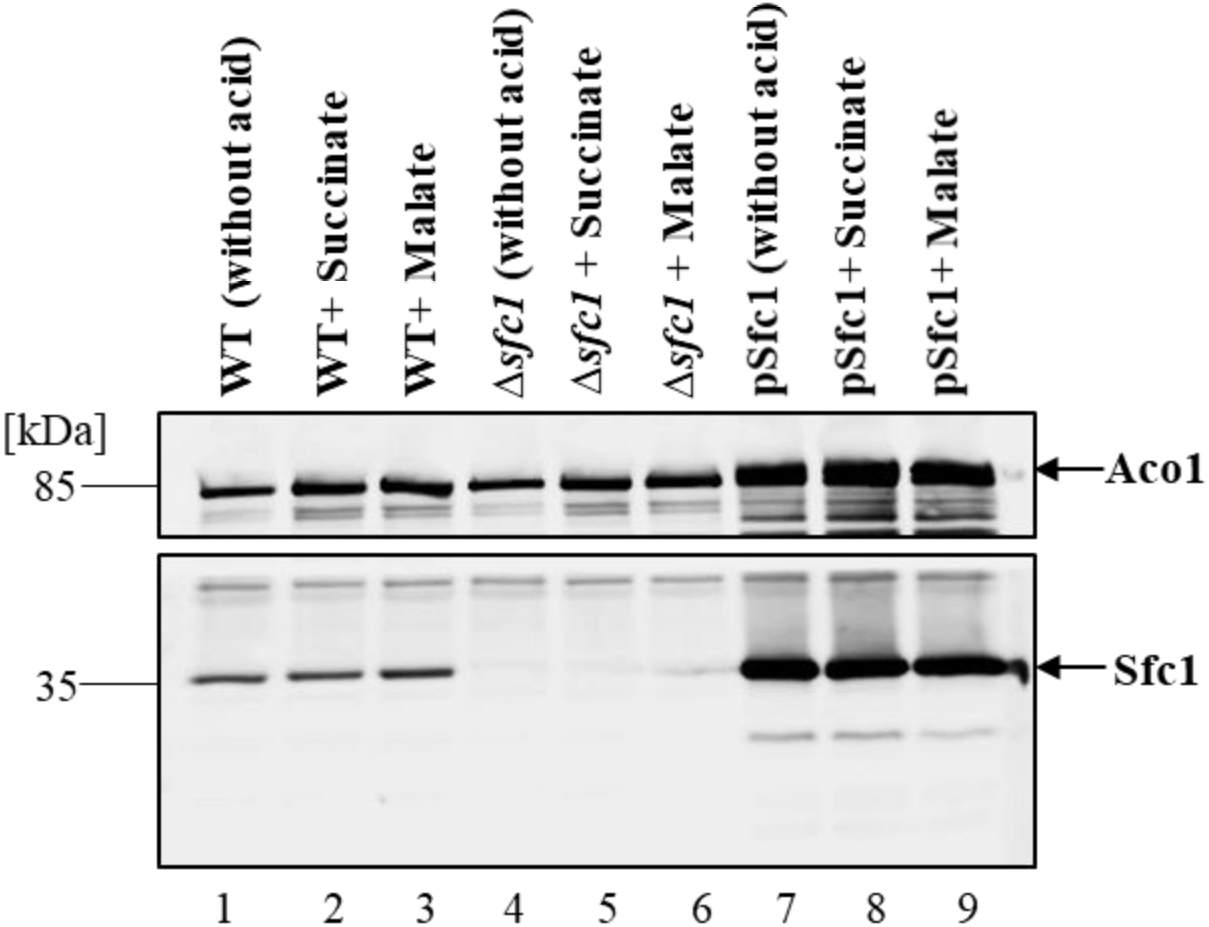
The expression levels of Sfc1 remain consistent, irrespective of the presence or absence of organic acid. Western blot analysis of WT, Δ*sfc1*, and pSfc1 in the absence (lanes 1, 4, and 7) or presence of succinate (lanes 2, 5, and 8) or presence of malate (lanes 3, 6, and 9) using anti-Sfc1 (bottom blot) and anti-Aco1 (top blot) antibodies. The upper band, corresponding to Aco1, was observed at approximately 85 kDa, while the lower bands correspond to Sfc1 at approximately 35 kDa.

**Figure S5.**
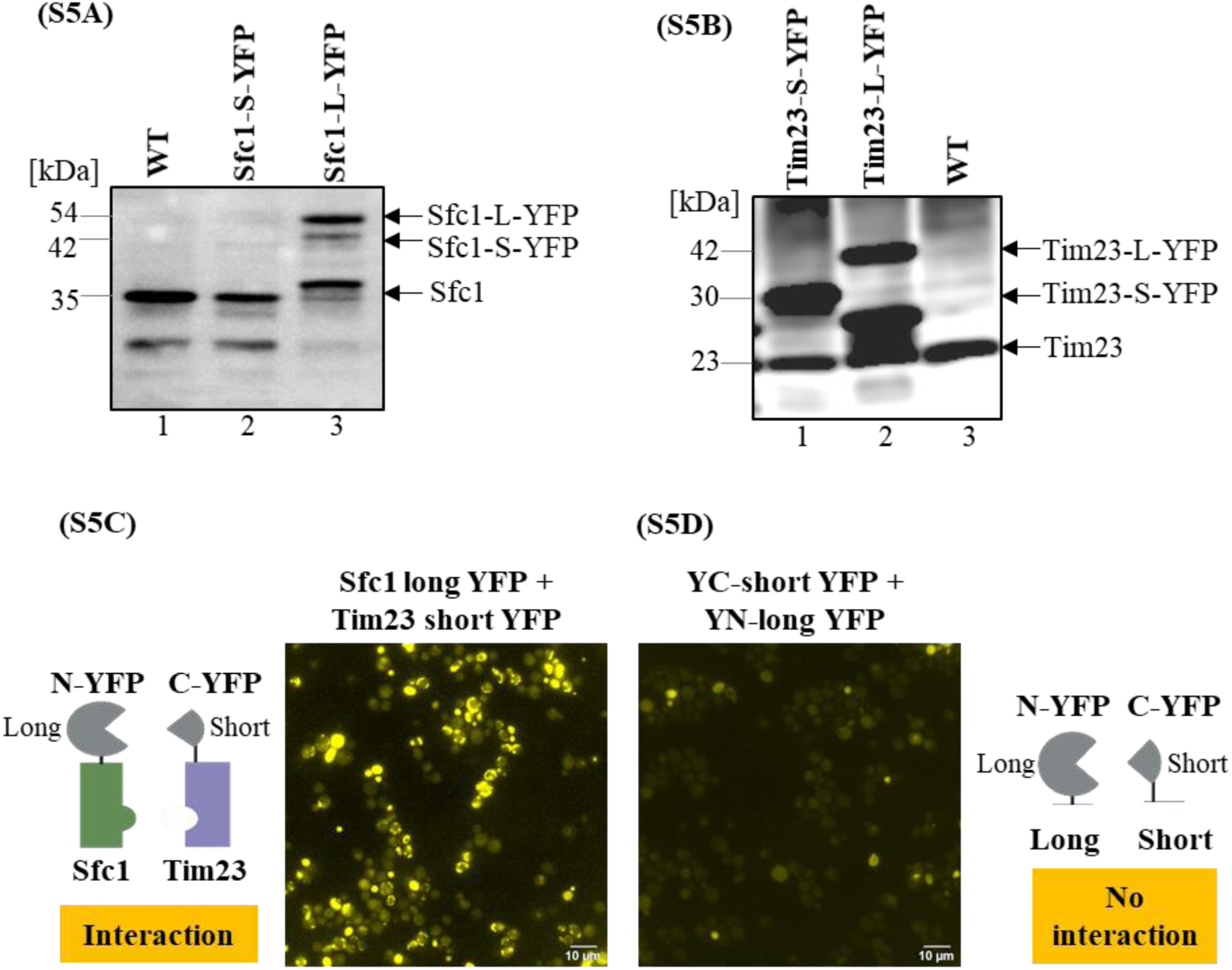
YFP expression in Sfc1 **(S5A)** and Tim23 **(S5B),** fused to long or short YFP was verified by western blot. **(S5A)** Lanes 1 and 3 correspond to wild-type (WT) and Sfc1 tagged with a long YFP, with sizes approximately at 35 kDa and 54 kDa, respectively. In contrast, Lane 2 correspond to Sfc1 tagged with a short YFP with (∼42 kDa), which was barely detectable, likely due to fragmentation of the short YFP. **(S5B)** Lanes 1, 2 and 3 correspond to Tim23 fused to short YFP, Tim23 fused to long YFP and WT Tim23 with sizes approximately 30 kDa, 42 kDa and 23 kDa respectively. **The interaction between Sfc1 and Tim23 is specific.** Yeast cells transformed with Sfc1 fused to long YFP and Tim23 fused to short YFP exhibit strong fluorescence **(S5C)**. In contrast, yeast cells transformed with a plasmid encoding both the short YFP and long YFP fragments yet without being attached to target proteins do not display fluorescence **(S5D).**

**Figure S6.**
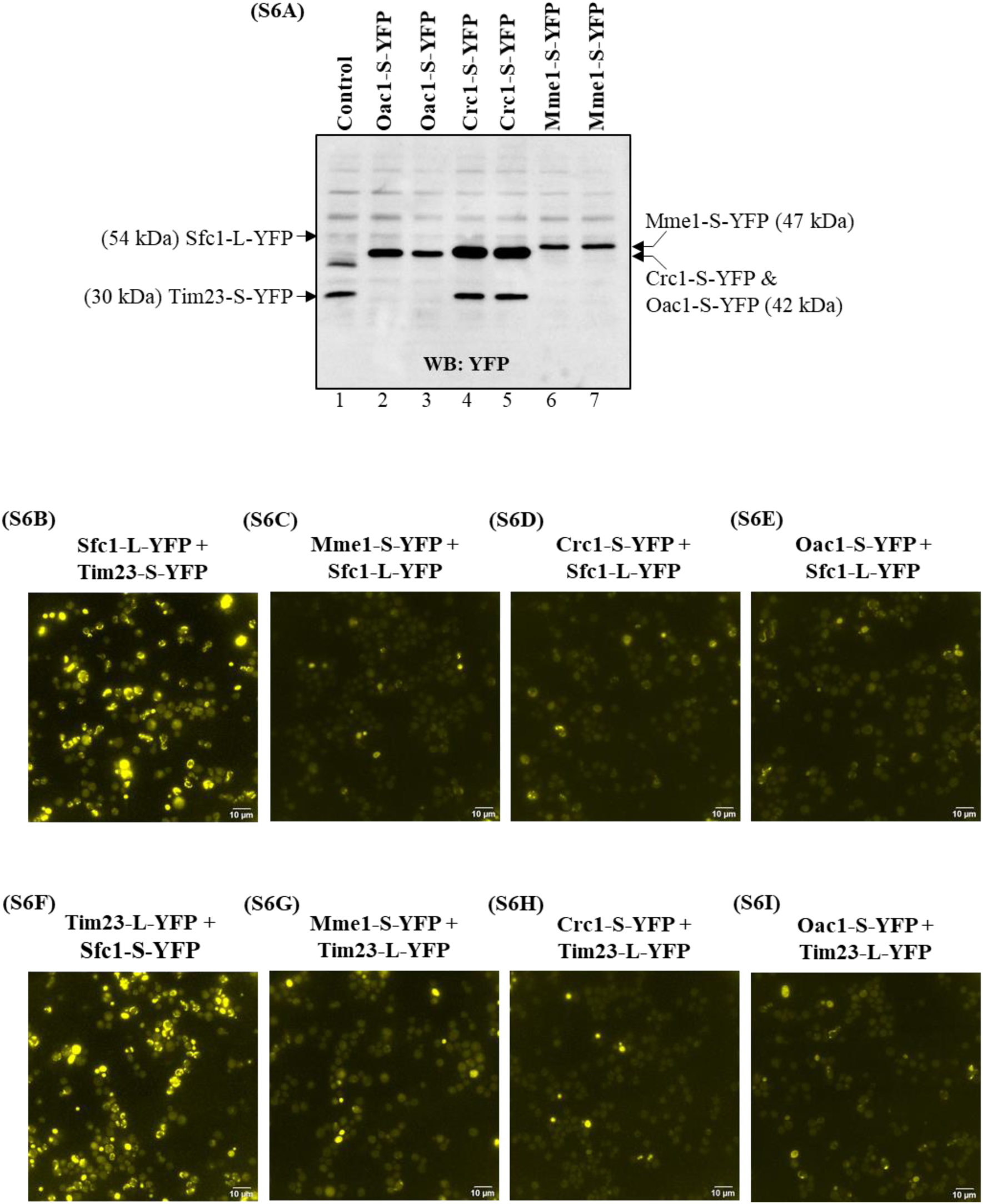
Other carriers do not behave like Sfc1. YFP expression in Mme1, Crc1 and Oac1 fused to short YFP were detected by western blot using anti-YFP **(S6A).** Lane 1 corresponds to control where Sfc1 is fused to long YFP (∼54 kDa) and Tim23 is fused to short YFP (∼30 kDa). Lanes 2, 3 correspond to Oac1 and lanes 4, 5 correspond to Crc1 fused to short YFP with molecular weight (MW) approximately 42 kDa and lanes 6, 7 correspond to Mme1 fused to short YFP with a MW approximately 47 kDa. As controls, yeast cells transformed with both Sfc1 fused to long YFP and Tim23 fused to short YFP, display a significant amount of fluorescence **(S6B)**, while yeast cells transformed with both Mme1 fused to short YFP and Sfc1 fused to long YFP do not show significant fluorescence **(S6C).** Similar results obtained with Crc1 **(S6D)** and Oac1 **(S6E).** In a parallel line of experiments, yeast cells were transformed with Mme1 fused to short YFP and Tim23 fused to long YFP, and again resulting in insignificant fluorescence in the cells (**S6G**) in contrast to the control **(S6F).** Similar results were obtained with Crc1 and Oac1 (**S6H** and **S6I**).

**Figure S7.**
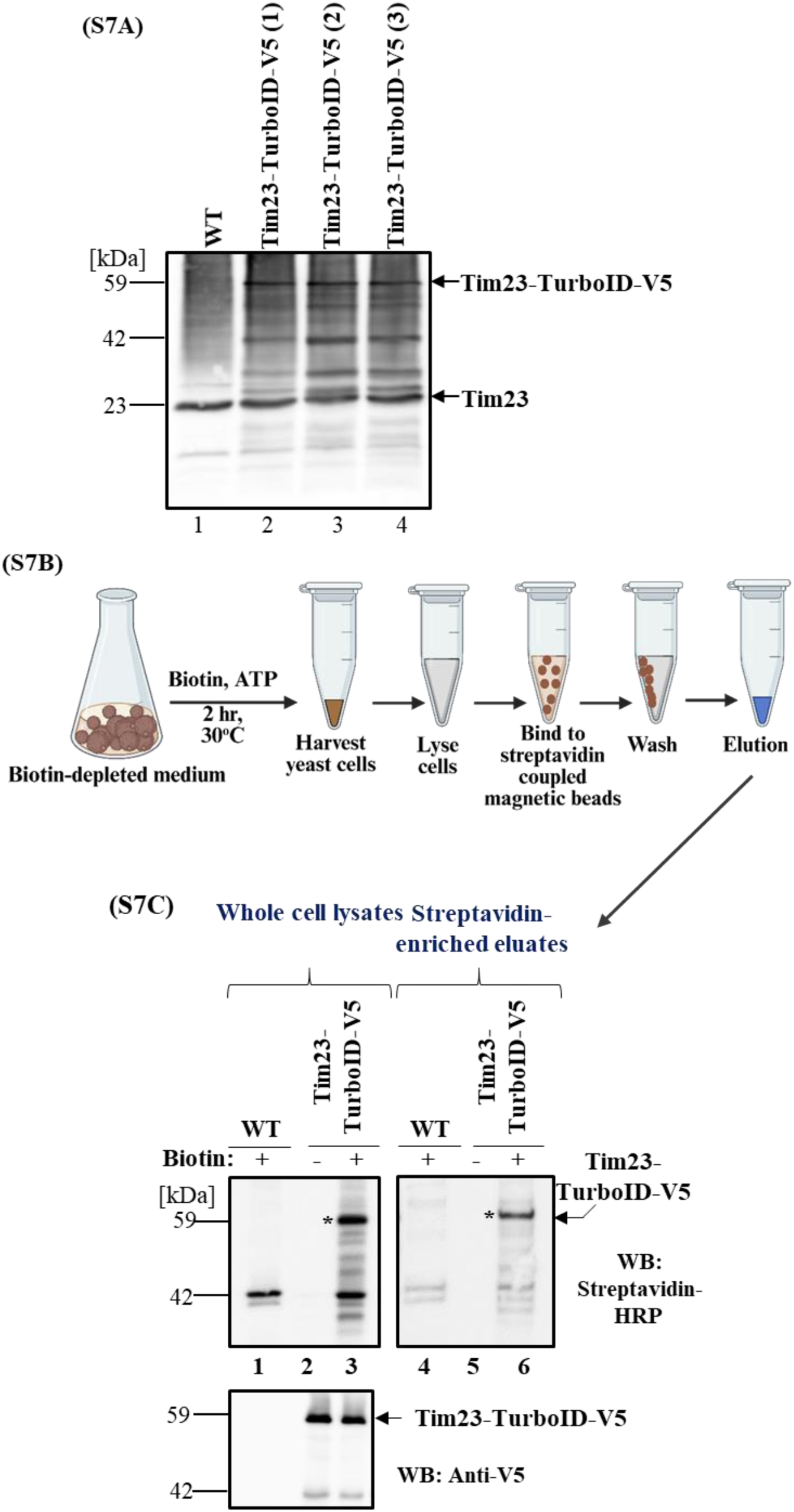
(S7A) TurboID identified Tim23 and Sfc1 as enriched proximate proteins. The expression of Tim23 fused to TurboID-V5 was confirmed by western blot (WB) using a Tim23 antibody. Lane 1 represents the control untagged strain, showing endogenous Tim23 at approximately 23 kDa. Lanes 2, 3, and 4 represent different clones of Tim23 fused to TurboIDV5, displaying a band at approximately 59 kDa (upper band) which is the fusion protein and a band at approximately 23 kDa (lower band) for endogenous Tim23. **(S7B) Overview of TurboID-based biotinylation approach in yeast.** Yeast cells were cultured in a biotin-depleted medium for 18 hours at 30°C (without biotin) and then supplemented with 50 µM biotin, 1mM ATP, and 5mM MgCl_2_ for an additional 2 hours at 30°C. The cells were lysed, and the biotinylated proteins were collected using streptavidin-coated beads and the biotinylated proteins were eluted and separated by SDS-PAGE. **(S7C)** WB of whole cell lysates from a control untagged strain (lane 1) and a Tim23-TurboID-V5 strain in the absence (lane 2) or in the presence (lane 3) of exogenous biotin are presented. Whole cell lysates were blotted with streptavidin-HRP antibody (top) to visualize biotinylated proteins and anti-V5 antibody (bottom panel) to detect ligase expression of TurboID. Lane 4 corresponds to streptavidin-enriched eluate of a control untagged strain and lanes 5 and 6 correspond to Tim23-TurboID-V5 in the absence and presence of exogenous biotin respectively. Asterisks indicate the position of the 59 kDa Tim23-TurboID-V5 protein, which is endogenously biotinylated protein in whole cell lysate (lane 3) and in streptavidin-enriched eluate (lane 6) respectively.

**Figure S8.**
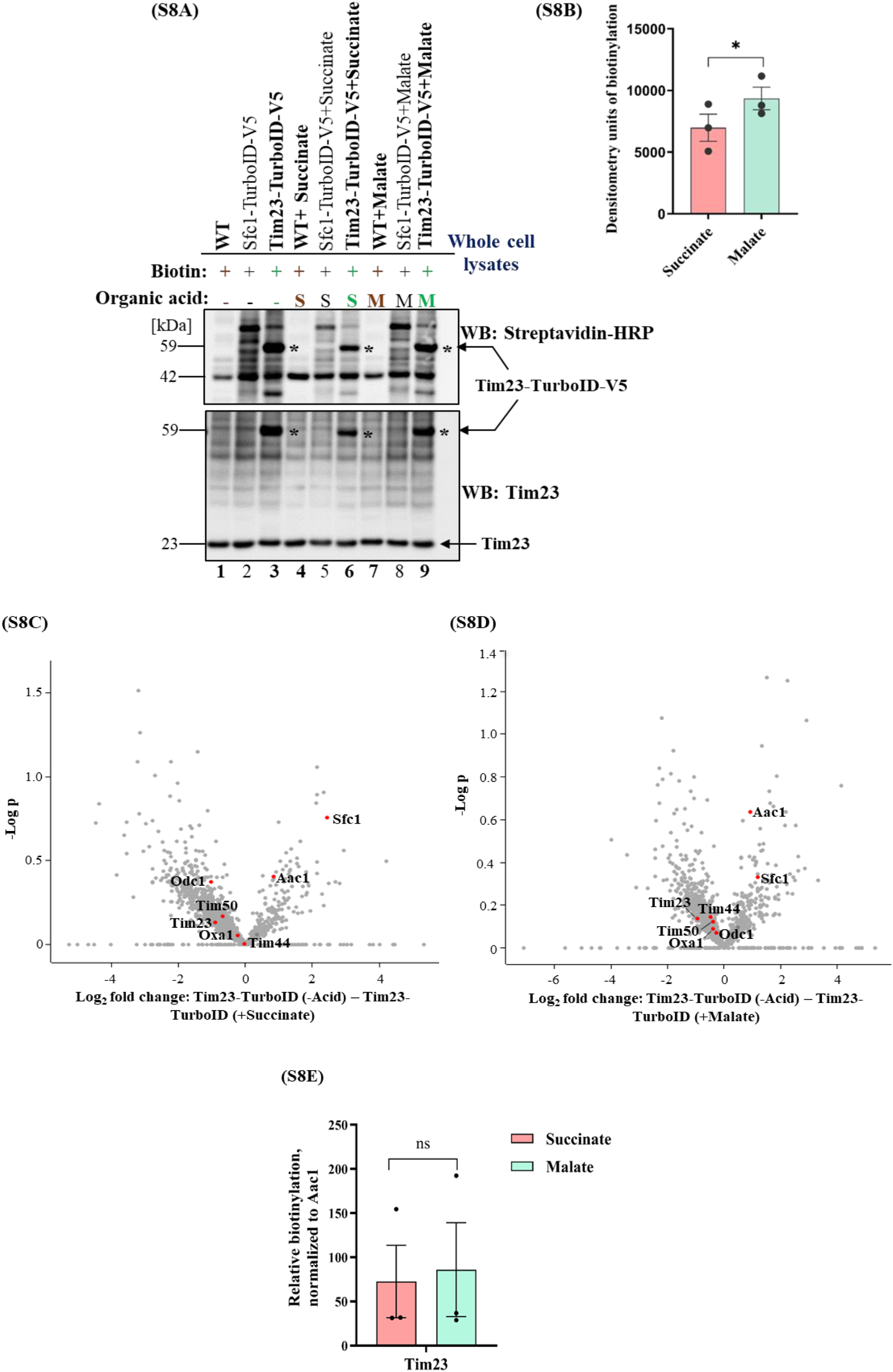
Succinate decreases the proximal biotinylation of Sfc1 by interfering with the interactions between Sfc1 and Tim23. **(S8A)** Western-blot analysis was performed using HRP-conjugated streptavidin (top) to detect biotinylated proteins and an anti-Tim23 antibody (bottom) to detect the endogenous expression of Tim23. The Tim23 protein fused to TurboID-V5 was expressed in strains grown in the presence of 50µM exogenous biotin without organic acid (lane 3), with 50mM succinate (lane 6), or with 50mM malate (lane 9). Asterisks mark the 59 kDa Tim23-TurboID-V5 fusion protein, which is endogenously biotinylated (top and bottom). In the wild-type untagged strain, no biotinylation of the target protein was detected (lanes 1, 4 and 7, upper gel). The anti-Tim23 blot also shows the endogenous Tim23 protein, approximately 23 kDa in size, observed across lanes 1-9. As a control for the Tim23-TurboID-V5 biotinylation reaction, we used Sfc1-TurboID-V5, which resulted in a distinctly different biotin labeling pattern (lanes 2, 5, and 8). **(S8B)** Subsequently, densitometry analysis of band intensities from the Tim23-TurboID-V5 strain grown in the presence of either succinate or malate was quantified and represented in a graph based on data from three independent experiments. The graph showed that the amount of biotinylation was reduced by approximately 37% in the presence of succinate compared to malate (mean ± SEM [n=3], two-tailed t-test *p=0.04). Volcano plots showing differential protein expression between control samples: without acid vs. succinate **(S8C)** and without acid vs. malate **(S8D)**. Fold change (log₂) is shown on the x-axis, and statistical significance (–log₁₀ p-value) on the y-axis. **(S8E)** Tim23-TurboID-V5 shows self-biotinylation, with a lower relative level observed in the presence of succinate compared to malate; however, this difference is not statistically significant.

**Figure S9.**
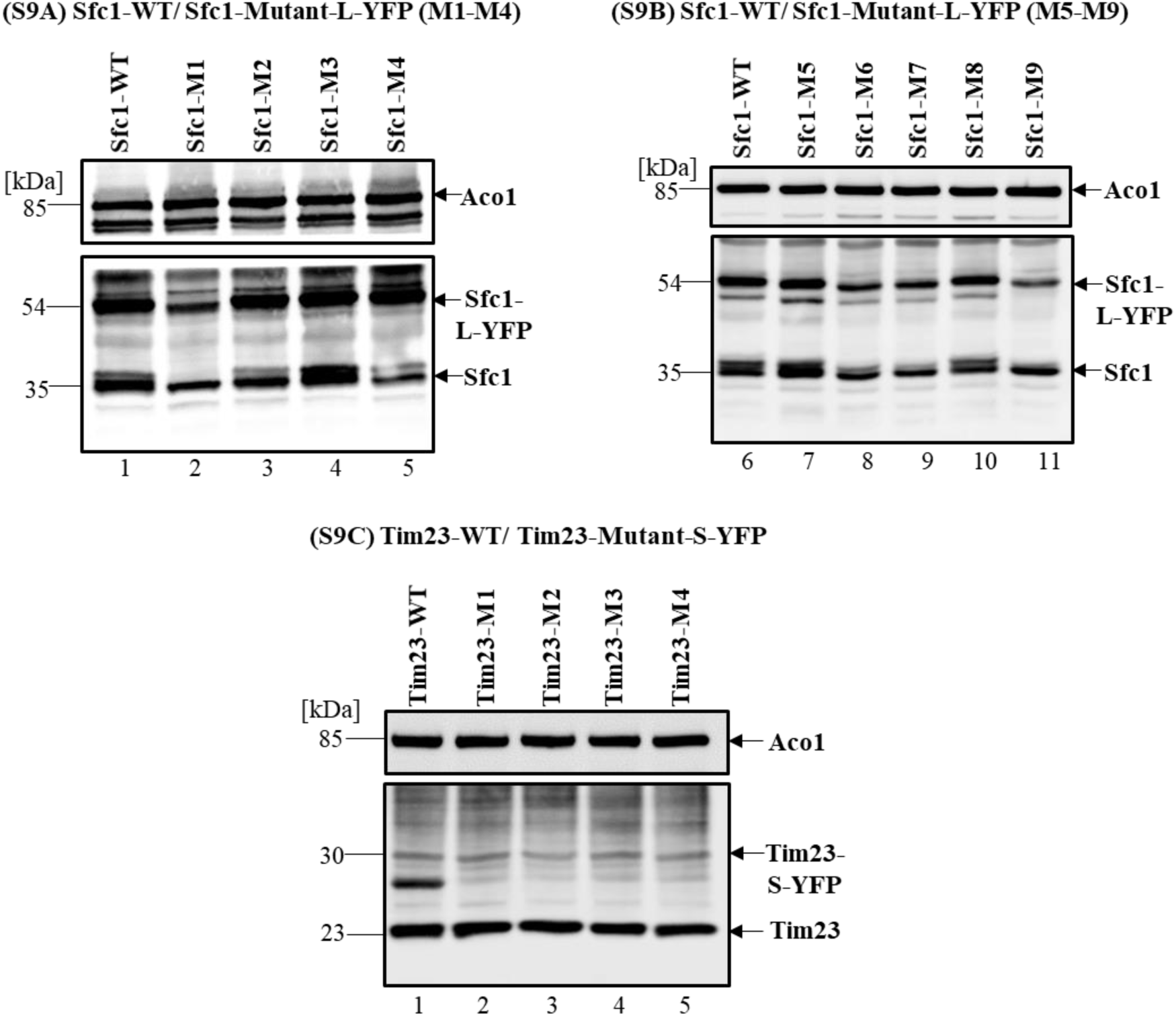
Expression of Sfc1 and Tim23 fusion mutants. **(S9A and S9B)** Western blot analysis confirmed the expression of Sfc1 mutant constructs fused to long YFP using an anti Sfc1 antibody (middle arrow) and an anti -Aco1 antibody as a control (top arrow, ∼85kD). Lane 1 and 6 represent the WT-Sfc1 strain fused to long YFP (bottom arrow), while lanes 2-5 and lanes 7-11 correspond to Sfc1-mutants tagged with long YFP. Arrows indicate the ∼35 kDa (endogenous Sfc1 protein) and ∼54 kDa (WT and Sfc1 mutant proteins with a long YFP); M: number of mutant. **(S9C)** Western blot analysis verified the expression of Tim23 mutant constructs fused to short YFP using an anti-Tim23 antibody (middle arrow, ∼30kD) whereas wild-type Tim23 (bottom arrow, ∼23kD) and aconitase (top arrow, ∼85kD) are controls. Lane 1, WT-Tim23 strain, while lanes 2–5 correspond to Tim23-mutants 1, 2, 3, and 4, fused to short YFP.

**Figure S10.**
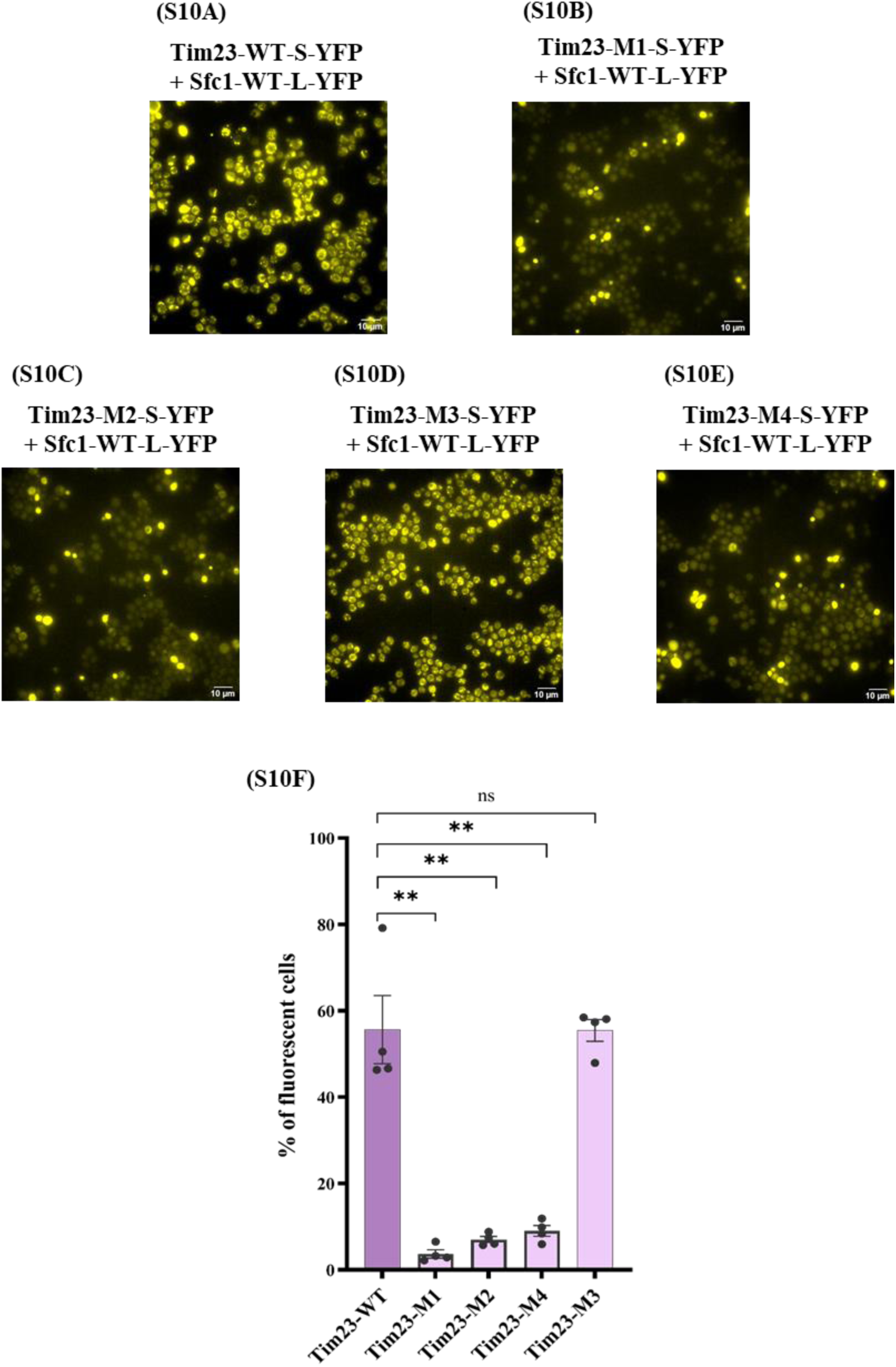
Alteration of the Tim23 sequence and its effect on the Tim23-Sfc1 interaction. Yeast strains were examined under the fluorescence microscope: WT-Sfc1 fused to long YFP and transformed with either **WT-Tim23** fused to short YFP **(S10A)**, **Tim23**-**M1** fused to short YFP **(S10B)**, **Tim23-M2** fused to short YFP **(S10C)**, **Tim23-M3** fused to short YFP **(S10D)** and **Tim23-M4** fused to short YFP **(S10E)**; M: number of mutant. Detailed information is provided in Supplementary Table 2. The graph quantifies the percentage of fluorescent cells relative to the total cell count **(S10F)**, with data derived from four independent experiments. (mean ± SEM [n=4], two-tailed t-test **p<0.01).

**Figure S11.**
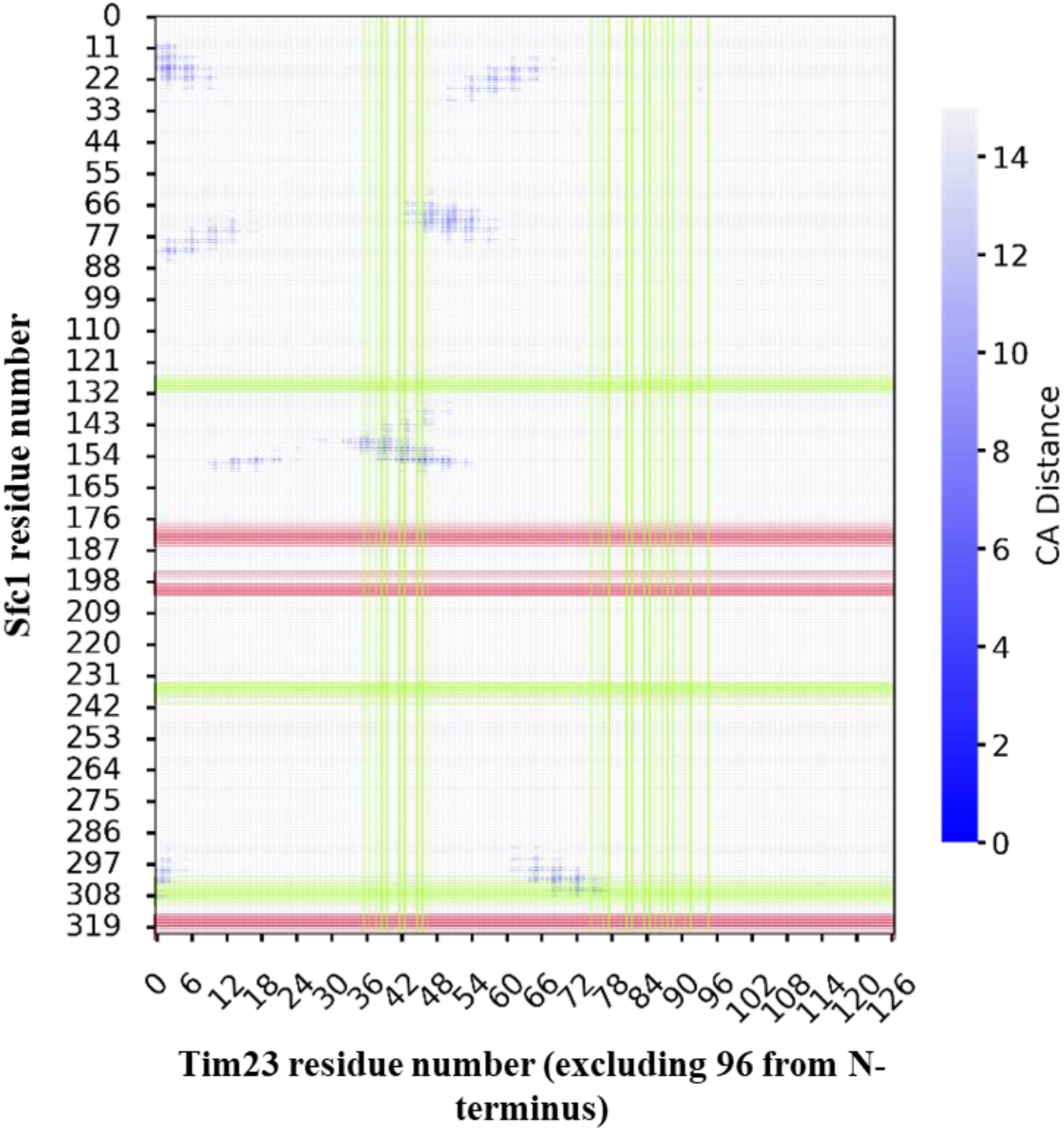
Interchain Distance-map. The interchain residue C*α* − C*α* distance (Å) matrix is shown here. Overlayed on the distance map is the fluorescence measure due to modifications in Sfc1 and Tim23. The residues that exhibit fluorescence post-modification are shown in green rectangles, and those that are non-fluorescent are shown in red rectangles. The residues at the interface, defined as those that are within 7 Å of each chain, are: **Sfc1:** Met15, Gly18, Thr19, Phe23, Leu26, Leu69, Tyr72, Leu75, Val79, Asn150, Gly152, Pro153, ASN156, Val300, His303. **Tim23:**Cys99, Tyr106, Lys132, Asn136, Leu139, Asn140, Ile142, Thr143, Lys144, Pro147, Phe148, Asn151, Ile155, Leu158, Ser165.

## Supplementary Tables

**Table S2.** Tim23 mutants used in the study. Different colors indicate mutant groups: **yellow** represents WT-like Tim23 fluorescence, while **purple** indicates decreased fluorescence in Tim23 mutants. (Δ=Deleted amino acid).

| *Name* | *Mutation description* | *% of fluorescent cells* |
| --- | --- | --- |
| *Tim23-WT* | - | 56 |
| *Tim23-Mutant-1* | ΔL132-K143 | 4 |
| *Tim23-Mutant-2* | ΔK172, ΔH173, ΔD174 | 7 |
| *Tim23-Mutant-3* | ΔE38, ΔH52, ΔD54, ΔR57 | 55 |
| *Tim23-Mutant-4* | ΔD12, ΔD13, ΔD22, ΔK25, ΔK27 | 9 |

**Table S3.** Interchain residues that are interacting. Interchain residues in the interface that are interacting with each other. The possible type of interaction is also mentioned here.

| **Sfc1** | **residue** | **Tim23** | **residue** | **Interaction** |
| --- | --- | --- | --- | --- |
| 303A | HIS | 98B | LEU | vanderwaals, electrostatic, hydrogen-bonding |
| 299A | TYR | 162B | ILE | hydrophobic |
| 303A | HIS | 97B | ASP | electrostatic, hydrogen-bonding |
| 23A | PHE | 158B | LEU | hydrophobic |
| 26A | LEU | 151B | ASN |  |
| 26A | LEU | 155B | ILE | hydrophobic |
| 19A | THR | 158B | LEU |  |
| 69A | LEU | 148B | PHE | hydrophobic |
| 296A | VAL | 158B | LEU | hydrophobic |
| 296A | VAL | 162B | ILE | hydrophobic |
| 300A | VAL | 99B | CYS |  |
| 23A | PHE | 155B | ILE | hydrophobic |
| 69A | LEU | 144B | LYS |  |
| 300A | VAL | 98B | LEU | hydrophobic |

**Additional Supplementary Tables will be provided at the time of publication.**

**Table S1: Tim23-specific interactors identified by TurboID.** Proteins specifically labeled by Tim23-TurboID compared with the untagged strain, identified by LC-MS/MS. Among ∼310 proteins, 37 were membrane proteins (18 mitochondrial).

**Table S4: List of plasmids used in this study.**

**Table S5: The primers listed below were used for cloning in this study.**

**Table S6: Plasmid combinations used for co-transformation in yeast for fluorescence microscopy.**

